# Creation of a single cell RNA seq meta-atlas to define human liver immune homeostasis

**DOI:** 10.1101/2021.02.12.430976

**Authors:** Brittany Rocque, Arianna Barbetta, Pranay Singh, Cameron Goldbeck, Doumet Georges Helou, Yong-Hwee Eddie Loh, Nolan Ung, Jerry Lee, Omid Akbari, Juliet Emamaullee

**Author notes:** **Corresponding Author:** Juliet Emamaullee MD PhD FRCSC FACS, 1510 San Pablo Street, Suite 412, Los Angeles, CA 90033.

## Abstract

The liver is unique in both its ability to maintain immune homeostasis and in its potential for immune tolerance following solid organ transplantation. Single-cell RNA sequencing (scRNA seq) is a powerful approach to generate highly dimensional transcriptome data to understand cellular phenotypes. However, when scRNA data is produced by different groups, with different data models, different standards, and samples processed in different ways, it can be challenging to draw meaningful conclusions from the aggregated data. The goal of this study was to establish a method to combine ‘human liver’ scRNA seq datasets by 1) characterizing the heterogeneity between studies and 2) using the meta-atlas to define the dominant phenotypes across immune cell subpopulations in healthy human liver. Publicly available scRNA seq data generated from liver samples obtained from a combined total of 17 patients and ∼32,000 cells were analyzed. Liver-specific immune cells (CD45+) were extracted from each dataset, and immune cell subpopulations (myeloid cells, NK and T cells, plasma cells, and B cells) were examined using dimensionality reduction (UMAP), differential gene expression, and ingenuity pathway analysis. All datasets co-clustered, but cell proportions differed between studies. Gene expression correlation demonstrated similarity across all studies, and canonical pathways that differed between datasets were related to cell stress and oxidative phosphorylation rather than immune-related function. Next, a meta-atlas was generated via data integration and compared against PBMC data to define gene signatures for each hepatic immune subpopulation. This analysis defined key features of hepatic immune homeostasis, with decreased expression across immunologic pathways and enhancement of pathways involved with cell death. This method for meta-analysis of scRNA seq data provides a novel approach to broadly define the features of human liver immune homeostasis. Specific pathways and cellular phenotypes described in this human liver immune meta-atlas provide a critical reference point for further study of immune mediated disease processes within the liver.

## INTRODUCTION

The liver is an immunologically complex organ with an abundance of resident leukocytes which comprise a significant proportion of residing nonparenchymal cells.(1,2) A unique but poorly understood feature of the liver is its physiologic immunotolerance, which is thought to be the result of the organ receiving much of its dual blood supply from the portal vein. The portal vein shuttles blood to the liver via the enterohepatic circulation, which carries both nutritionally-derived and bacterial antigens without causing an untoward inflammatory response.(3) Further characterization of the unique hepatic immune microenvironment has the potential to aid study of a spectrum of immune-mediated disease processes of the liver including autoimmune hepatitis, cholangiopathy, transplant rejection, and the tumor microenvironment of intrahepatic malignancy.(4)

Single cell RNA sequencing (scRNA seq) has led to rapid advances in our understanding of cellular phenotypes of both *in situ* human tissues in recent years.(5,6) Application of scRNA seq to human liver tissue to examine both healthy and pathogenic states at the molecular level has proven useful to identify heterogeneity among various cell types, including immune cell populations and epithelial progenitors.(7–9) Generation of these libraries through high-throughput sequencing techniques such as 10x Genomics or Drop-seq generates highly dimensional data, which is suitable to both answer and also to rapidly generate hypotheses. Given the relatively high cost of scRNA seq, a typical human study only analyzes a small number of unique samples (as few as three unique patients), despite the known genetic heterogeneity across individuals.(10,11) In 2018, the NIH updated the policy for data access of genomic datasets, with the goal of enhancing researchers’ ability to perform pooled analyses or meta-analyses on genomics data particularly in human specimens.(12) More recently, there are several NIH requests for applications specifically targeted at secondary and integrated analytic approaches to harness the power of these existing, funded datasets.(13)

Combining liver-specific immune cell subsets of scRNA seq datasets yields a higher number of cell libraries across a larger sample of patients, which is crucial when considering that knowledge generated from these human “atlases” may be used to understand biochemical mechanisms of disease and to develop therapeutics for clinical use. One major drawback of combining unique studies is that differences in technique have been demonstrated to lead to unexpected alterations in sequencing results.(14,15) A study by Bonnycastle *et al* used scRNA seq of human pancreatic islet cells and compared different tissue processing techniques, including fresh, fixed, and cryopreserved tissues. Despite processing tissue samples from the same source, this analysis demonstrated differences among cell type proportions recovered as well as changes in gene expression signatures across different processing techniques.(16) Thus, combining multiple human liver scRNA datasets has the potential to account for more biological variability across different patients, potentially attenuating the effects of pre-analytical variables that are not biologically meaningful.

The aim of this study was to determine if a meta-analysis of existing normal human liver scRNA seq datasets could be performed to generate a comprehensive human liver immune meta-atlas for future use as a reference point for examining immune-mediated liver diseases. In Part 1, an extensive comparison was performed between studies in order to establish whether generalizability across datasets would be meaningful. In Part 2, the pooled meta-atlas was explored to define expression profiles across four major immune subpopulations and to provide a re-usable meta-atlas of healthy liver immune homeostasis.

## RESULTS

### Part 1: Approach to meta-analysis of liver immune scRNASeq data

To begin, immune-specific subsets of human liver tissue were identified and extracted from three unique studies: “LACE^e^”, “L^nb^” and “LCD45^e^” (**Figure 1**).(7–9) Techniques used for the RNA sequencing of individual datasets are summarized in **Table 1**. The combined dataset included single-cell RNA sequencing of 17 normal human liver samples with approximately 32,000 hepatic CD45+ cells.

**Table 1.** Comparison of methods between three peer-reviewed single cell RNA sequencing studies of the liver.

|  | LACE<sup>e</sup> | L<sup>nb</sup> | LCD45<sup>e</sup> |
| --- | --- | --- | --- |
| **Samples** | 9 liver resection pts mCRC ICC | 5 caudate lobes of DBD Ltxp donors | 3 adult Ltxp donors blood, spleen, liver |
| **Perfusate** | HEPES | HTK solution, then HBS + EGTA | Not addressed |
| **Cell fractionation** | PHHs and NPCs isolated, mixed then FACS | None | Cell filtration, centrifugation and Ficol |
| **Removal of non-viable cells** | Gradient centrifugation | Trypan blue exclusion | eFluor 450 exclusion |
| **Preservation method** | Mix of cryopreserved and fresh | Fresh | Fresh |
| **Cell enrichment** | Yes - FACS | None | Yes - FACS |
| **Enrichment for Lymphocytes** | No | No | Yes - marker CD45 |
| **Enrichment for LSECs** | Yes - marker CLEC4G | No | No |
| **Enrichment for MaVECs** | Yes - marker CD34, PECAM | No | No |
| **Enrichment for Cholangiocytes** | Yes - marker EPCAM | No | No |
| **Removal of low-quality cells** | Yes - excluded >2% KCNQ1OT1 transcripts | Removed >50% mitochondrial content | No |
| **Number of cells scRNA seq** | 10,372 | 8,444 | 70,706 |
| **scRNA seq technique** | mCEL-Seq2 | 10x | 10x |
| **Identification of cell types** | RaceID3 (mintotal = 1000, minexpr = 2, minnumber = 10, outminc = 2, cln = 15) | Not addressed | Seurat v3 |
| **Samples across patients** | Co-clustered | Co-clustered | Co-clustered |
| **Different preps for patients** | Co-clustered | Not applicable | Not applicable |
| **Cell doublets** | Not addressed | Not removed | Removed |

LACE<sup>e</sup> Aizarani et al. A human liver cell atlas reveals heterogeneity and epithelial progenitors
L<sup>nb</sup> MacParland et al. Single cell RNA sequencing of human liver reveals distinct intrahepatic macrophage populations
LCD45<sup>e</sup> Zhao et al. Single-cell RNA sequencing reveals the heterogeneity of liver-resident immune cells in human

**Figure 1:**
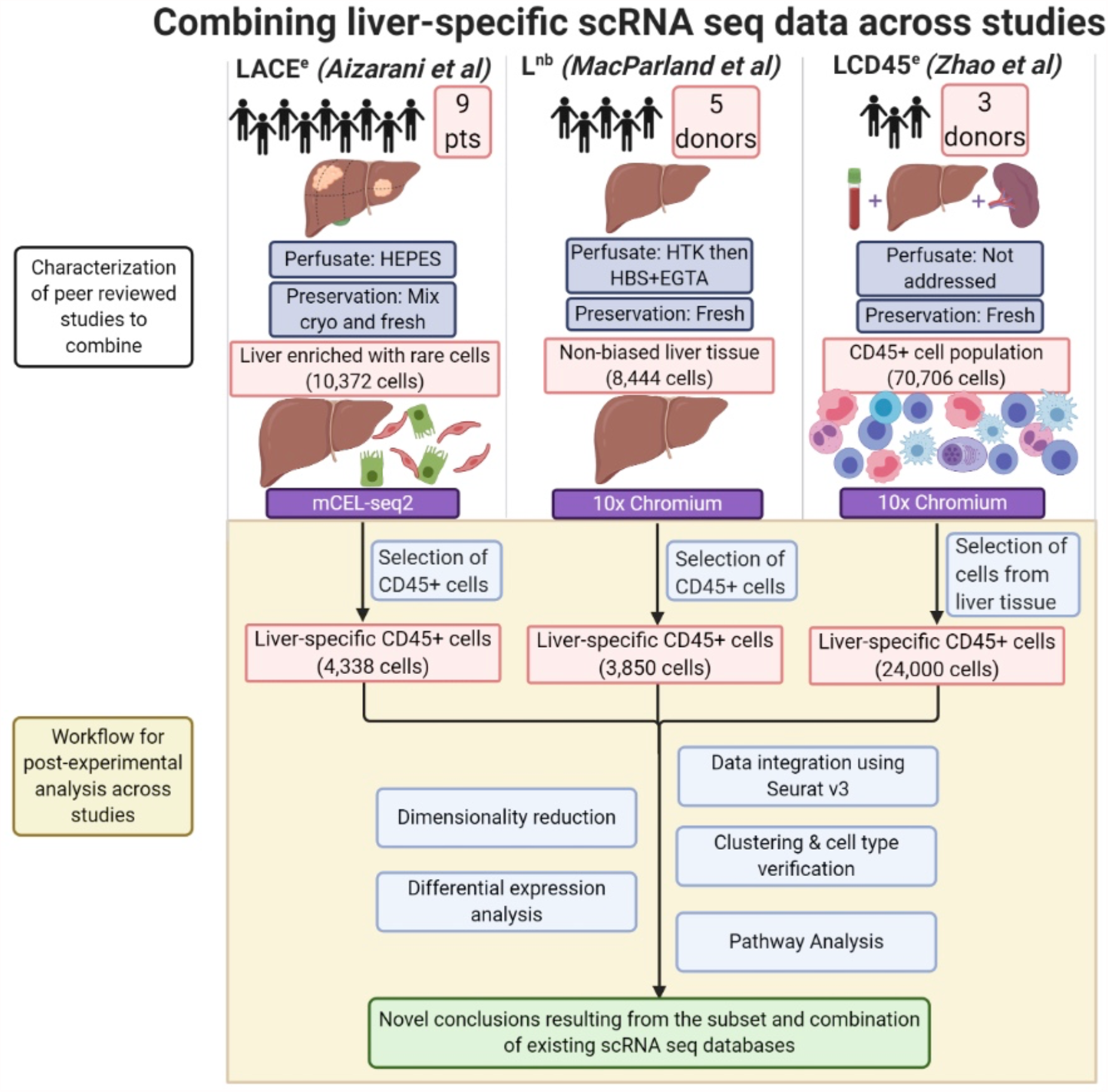
Combining liver-specific scRNA seq data to create a human liver immune meta-atlas. Schematic diagram summarizing the number of subjects and major pre-analytical variables pertaining to scRNA sequencing of human liver. Post-experimental analysis performed in this study is highlighted in the shaded region. CD45+ cells were selected from the LACE^e^ and L^nb^ studies. CD45+ cells (which were already pre-selected for in the LCD45^e^ dataset) which were of liver origin were post-experimentally extracted as splenic and peripheral blood cells were also included in the original dataset. The scRNA seq analysis pipeline (Seurat v3 in R) was applied, then clustering and differential expression analysis were used to assess the ability to combine scRNA seq techniques and generate datasets for tertiary pathway analysis of immune function in the normal human liver. Created with BioRender.com.

Clustering analysis was performed on all three datasets individually to detect differences in global cellular phenotypes (**Figure 2A-C**). Projection of all three datasets to the same UMAP coordinates revealed a homogeneous interdigitation of each cluster, however the LCD45^e^ subset comprised the majority of cells (∼24,000 cells compared to ∼4,000 cells in each of the other two studies; **Figure 2D**). Pairwise gene expression correlation analysis was performed which revealed that the LACE^e^ dataset had less concordance with both the L^nb^ (*R*=0.81; **Figure 2E**), and LCD45^e^ datasets (*R*=0.79; **Figure 2F**). L^nb^ and LCD45^e^, both obtained using the 10x platform, demonstrated a more idealized relationship (*R*=0.95; **Figure 2G**).

**Figure 2:**
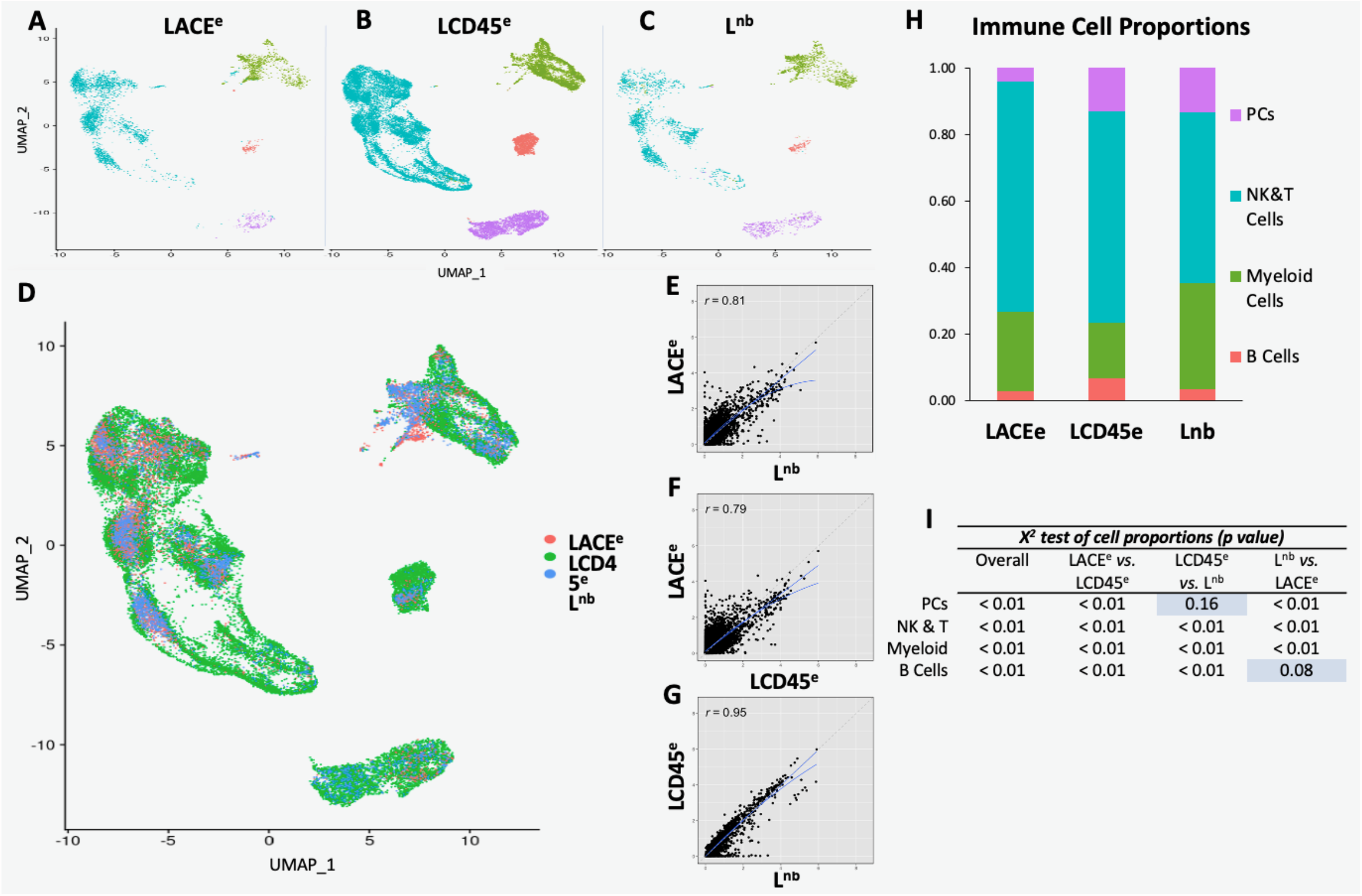
Data visualization with clustering and gene expression correlation between datasets. **(A-C)** UMAP plots of individual studies. **(D)** Plot showing co-clustering of three single cell datasets. **(E-G)** Gene expression correlation plots with solid lines showing linear and quadratic regressions. Dashed line representing idealized relationship. **(H)** Bar graph representation of immune cell proportions of each individual dataset. **(I)** *X*^2^ comparison showing significant differences in proportions of immune cell subpopulations. With pairwise analysis, most chi-squared values reached statistical significance except for plasma cells in LCD45^e^ versus L^nb^ and for B Cells in L^nb^ versus LACE^e^ (indicated by blue rectangles).

### Characterization of the frequency and proportion of immune cell subpopulations between liver scRNA seq datasets

Next, leukocyte subpopulations were identified based on expression of major cell-specific lineage marker genes and compared between the individual datasets (**Supplemental Figure 2**). Clusters were then designated as NK and T cells (combined), myeloid cells, B cells and plasma cells, representing the four major groups of interest for our study.

When compared between all three datasets there were differences in proportions of each immune cell subpopulation (*X*^*2*^ test, *p*<0.01 for each cell type; **Figure 2I**). Pairwise analysis between datasets also revealed differences in cell-type composition, and only two conditions did not reach statistical significance: comparison of LCD45^e^ and L^nb^ in plasma cells (*X*^*2*^ test, *p*=0.16) and comparison of L^nb^ and LACE^e^ in B cells (*X*^*2*^ test, *p*=0.08), which may be a consequence of B cells and plasma cells representing the smallest populations of cells between datasets. This observed scarcity of liver resident B cells and plasma cells is consistent previous reports of liver immune cell composition.(2) Despite some differences in cell-type recovery, the ranking of abundance of each cell type was preserved between datasets, such that NK and T cells were the most numerous (ranging from 51%-69% of cells), followed by myeloid cells (18%-32%), plasma cells (4%-14%), and then B cells (2%-7%; **Figure 2H**).

### Comparative analysis of leukocyte subpopulations between datasets reveals concordance, despite differences in liver tissue handling and sequencing techniques

A deeper analysis of gene expression profiles within immune cell subpopulation was conducted to identify differences in cellular phenotypes. Analysis of housekeeping genes was conducted, as an internal control of biological noise, thus serving as a marker of technical noise between datasets.(28) Expression analysis was performed on an established set of human housekeeping genes across immune cell subpopulations and between datasets using log normalized expression values (**Supplemental Figure 3**). Despite a relatively good agreement between means, comparison with one-way ANOVA identified differences in housekeeping gene expression (*p*<0.01). Pairwise analysis also showed significant differences in expression levels, which is likely the result of the large number of cells included and is thus a highly powered test.

Differences in gene expression profiles across immune cell subpopulations are anticipated due to their distinct biological functions. The number of differentially expressed (DE) genes was quantified between individual immune cell subpopulations (**Supplemental Figure 4**). Volcano plots were used to illustrate meaningfully DE genes, which were plotted as blue dots and indicate a fold-change ratio in expression of either <0.8 or >1.25 and Bonferroni corrected *p*-value<0.05. Non-DE genes were plotted as red dots. B cells had the fewest DE genes between all datasets, whereas cells of the myeloid lineage had the most DE genes. No individual dataset was an outlier with respect to the quantification of differential expression, but the LCD45^e^ dataset had the highest number of DE genes in B cells and myeloid cells, and the lowest number in plasma cells and NK and T cells.

Due to heterogeneous gene expression between leukocyte subpopulations (**Supplemental Figure 4**) and differences in cell proportions (**Figure 2H,I**), both of which could potentially bias dataset correlation analysis, pairwise expression correlation between datasets was performed and stratified by leukocyte subpopulation (**Supplemental Figure 5A-D and Supplemental Figure 6**). Comparison of L^nb^ and LCD45^e^ showed a more idealized relationship (*R*=0.93-0.96 depending on leukocyte subpopulation, **Supplemental Figure 5A.vi.-D.vi**), while LACE^e^ correlated less with the other two studies (**Supplemental Figure 5A.iv**,**v-D.iv**,**v**). The pattern of gene expression correlation was also analyzed using Rank-Rank Hypergeometric Overlap (RRHO) heatmaps (**Supplemental Figure 5A-D, panels i-iii**).(29) This technique uses heatmaps to indicate the degree of ranked differential expression agreement, where yellow indicates strong statistical overlap and blue represents no statistical overlap between corresponding axes. These panels represent overlap of differential expression in ranked form such that the bottom left of the panel represents the most upregulated genes, and genes in the upper right portion of the panel represent the downregulated genes (thus a higher rank corresponds to downregulation). When comparing NK and T cell expression patterns in LACE^e^ versus L^nb^, there are two heatmap regions with a high degree of statistical overlap (**Supplemental Figure 5A**). Examination of the remaining two pairwise comparisons in NK and T cells from different datasets, the overlap among the more highly expressed genes is evident, but genes at lower expression levels have less overlap. Plasma cells and B cells also exhibit RRHO agreement between studies. For myeloid cells, LACE^e^ and L^nb^ have better RRHO for the most highly expressed genes, whereas L^nb^ and LCD45^e^ have better agreement in genes expressed at lower levels. There is a bimodal agreement between LACE^e^ and LCD45^e^ (**Supplemental Figure 5B**).

### Differential gene expression in leukocyte subpopulations highlights the degree of agreement between datasets

The RRHO analysis demonstrated that there was relatively strong correlation with genes at the highest levels of expression. To begin to identify the dominant phenotype for each leukocyte subpopulation, the top 100 most abundant transcripts were identified for each cell type in L^nb^ which was the dataset with the most agreement with the other two datasets based on correlation (**Figure 2E-G**). A comparison was performed to determine if the same genes were present as the top 100 most abundant transcripts for each immune cell subpopulation of the other two studies. Depending on leukocyte subpopulation, LCD45^e^ had between 87 and 94 out of 100 ‘top genes’ in agreement. Alternately, LACE^e^ had rather poor agreement with top matching genes for each leukocyte subpopulation (**Supplemental Figure 7**).

Analysis was then performed between datasets in order to quantify how many genes were DE in each immune cell subpopulation (**Figure 3B**). In this analysis, DE genes with a fold change ratio <0.8 or >1.25 and a Bonferroni corrected p-value <0.05 are represented in volcano plots and compare individual gene expression level within an immune cell subpopulation, between datasets.(16) Some comparisons yielded a high number of DE genes between datasets (e.g., 3526 genes when comparing LCD45^e^ and LACE^e^ in myeloid cells). In contrast, when comparing the L^nb^ dataset to the LCD45^e^ dataset among the B cell population, there were only 767 DE genes and the myeloid cell population between these two datasets had 924 DE genes, again demonstrating fewer expression differences between L^nb^ and LCD45^e^ relative to LACE^e^.

**Figure 3:**
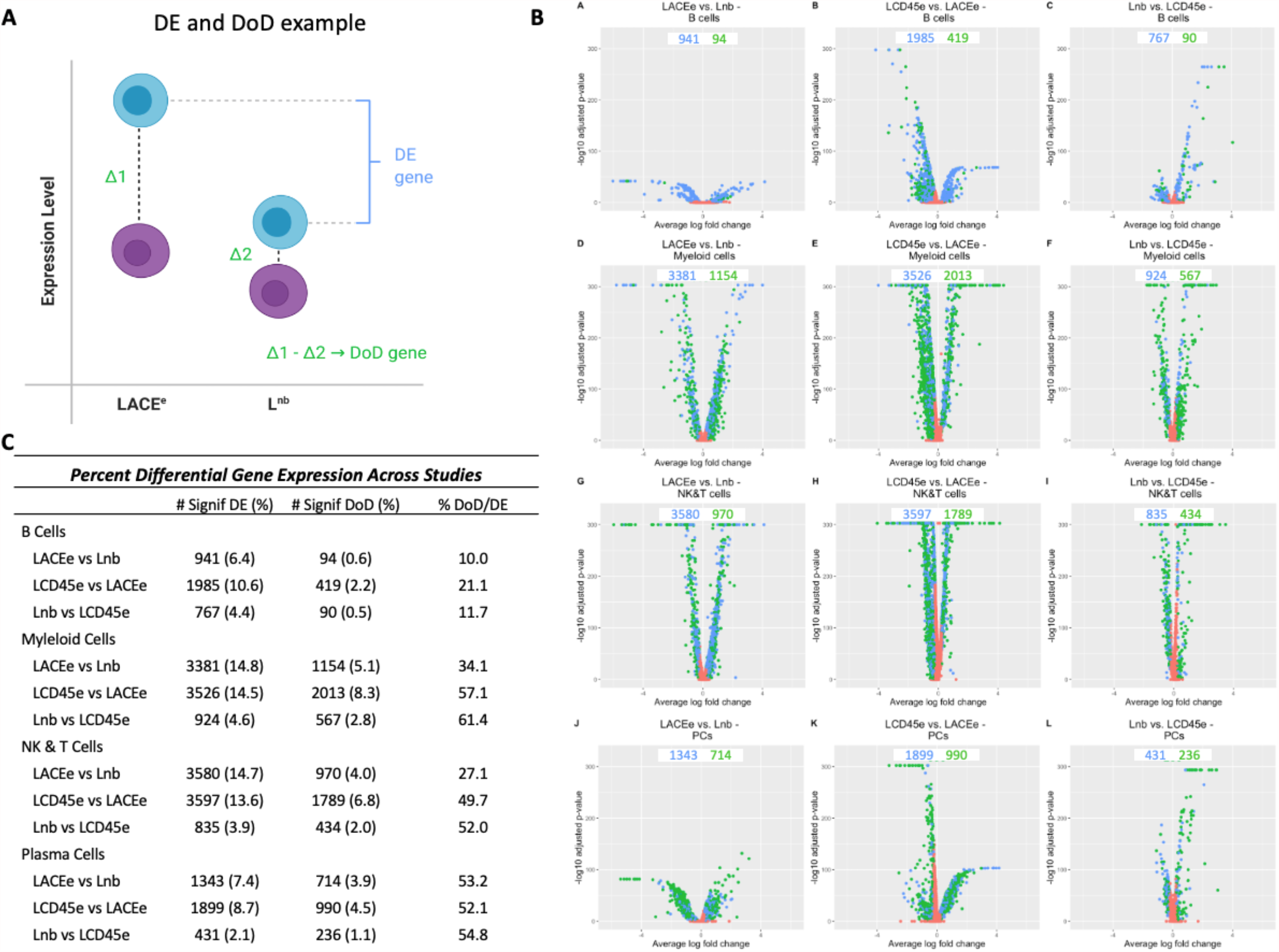
Differential gene expression analysis between datasets. **(A)** Schematic diagram for identification of a differentially expressed (DE) gene. The blue cells represent different expression levels for a gene in each leukocyte subpopulation (e.g., expression in B cells) and the purple cells represent the expression levels of that same gene, but in all other cell subpopulations. Differential expression only takes levels from the B cell into account. We also examine the Difference of Differences (DoD) which takes the difference in expression between B cells and all other cell types for dataset #1 and subtracts it from the differences in expressed between B cells and all other cell types for dataset #2 which may help categorize the differential expression that is accounted for by technical differences between studies (such as sequencing depth) rather than biologically relevant changes. **(B)** Volcano plots showing the number of DE genes (blue dots) and DoD genes (green dots) for each dataset condition as pairwise comparisons between studies and stratified by cell type. **(C)** Lists of differential gene expression numbers, DoD genes and then the percent of DoD genes out of the number of DE genes.

To better characterize the heterogeneity in gene expression, the difference of differences (DoD) was calculated between datasets (pairwise) and stratified by leukocyte subpopulation (particular versus all other). This metric involved calculating a gene’s expression difference between the cell subpopulation of interest and all other cell types and then comparing that difference to the difference seen in another dataset (**Figure 3A**). For example, a B cell in the LACE^e^ dataset has a difference of expression of the gene CD25 compared to all other cell types (represented as Δ1), the difference in CD25 expression between B cells and other cell types in L^nb^ is Δ2. Subtracting Δ2 from Δ1 yields the DoD, (**Figure 3**) which we used to approximate the proportion of DE genes where expression alteration could be explained by changes relative to other immune cell subpopulations (DoD significance defined as: DE with a fold change ratio <0.8 or >1.25 with *P*-adj_Bonfcorr_<0.05 and DoD *P*-adj_Bonfcorr_<0.05). We also anticipate that due to comparison between datasets and across differing technical platforms (10x Genomics versus mCel-seq) that there will be differences in the depth of sequencing, a phenomenon which has been known to affect differential expression analysis when comparing scRNA seq datasets.(30) Given this, we propose that our DoD analysis “explains”, or at least accounts for, some of the differential expression. Figure 3C lists the number of DE and DoD genes for each analysis as well as a percent total. The last column in the table shows the proportion of DE genes that also had a significant DoD (the ratio of the two previous columns). Notably in comparing myeloid cells between LACE^e^ and L^nb^, there are a high number of DE genes, but only 34% of these are also DoD genes. This suggests that nearly 70% of the observed differences of myeloid cells between these two datasets can be accounted for by comparing to the average expression level in the remaining cells of each rather than representing a unique feature of the myeloid population.

### Dominant gene expression profiles for immune cell subpopulations are concordant between datasets

Once DE genes were identified, the most significantly DE genes were chosen among each dataset to determine if an integrated dataset could identify dominant gene expression signatures associated with each immune cell subpopulation within human liver. The topmost DE genes for each leukocyte subpopulation in each individual dataset were identified. All three datasets’ candidate genes were combined to create a master list of top DE genes for each immune cell subpopulation (**Figure 4D**). Expression levels were plotted as a heatmap for each individual cell and organized by immune cell subpopulation (**Figure 4A-C**). Some heterogeneity was expected within cell type, as the myeloid and NK and T cell populations represent a variety of unique subsets. Despite this, the heatmaps demonstrate that candidate genes listed are indeed DE in the immune subpopulation of interest when compared with all other subpopulations and represent reproducible markers between each dataset, regardless of processing technique. The expression patterns outlined in the heatmaps have excellent agreement across studies, confirming that these datasets could be integrated to produce a meta-atlas of human liver immune function.

**Figure 4:**
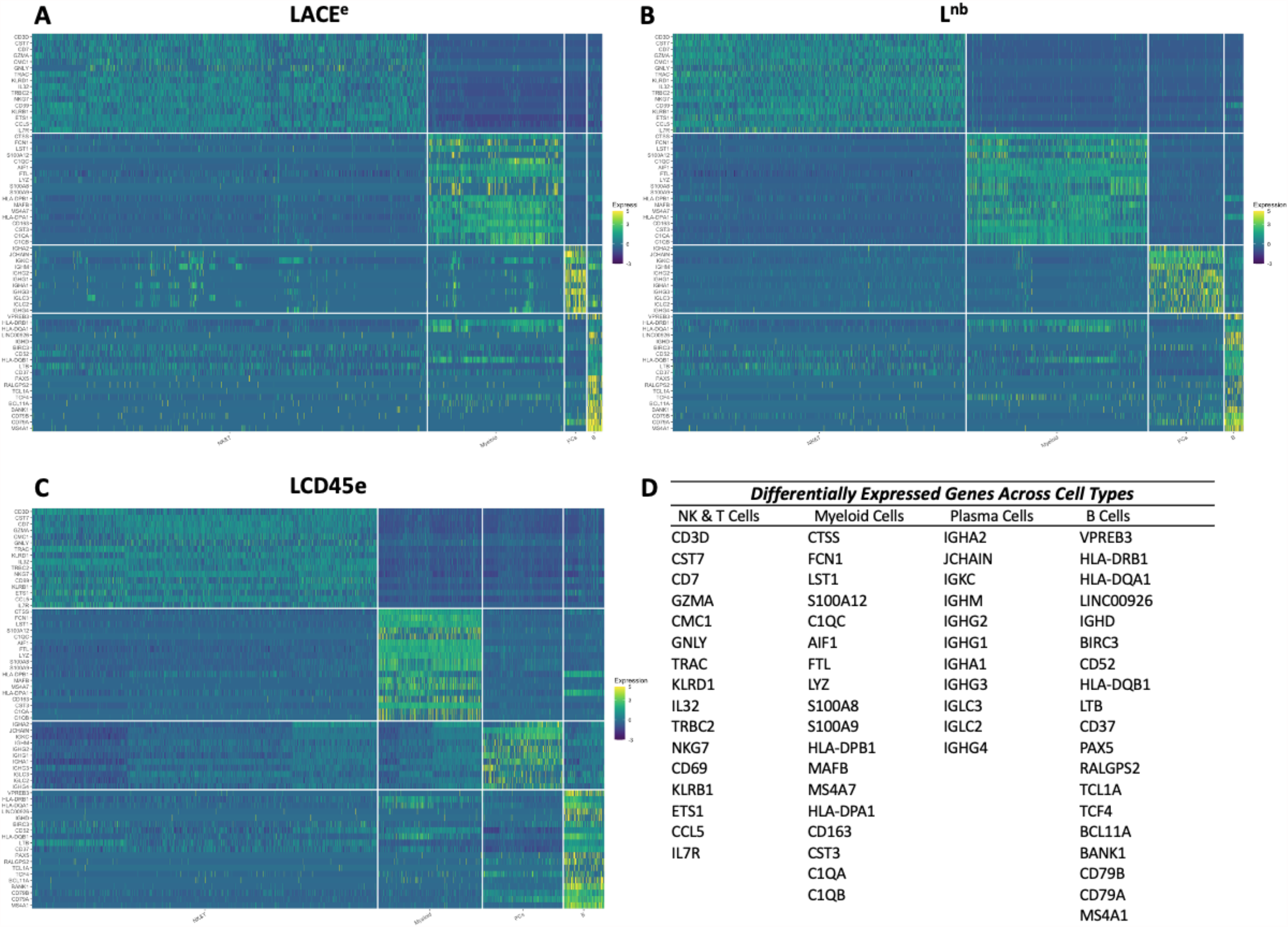
Heatmap representation of dominant differentially expressed genes between each immune cell subpopulation in each dataset. **(A)** Heatmap of the most DE genes across leukocyte subpopulation plotted using cells from the LACE^e^ dataset. **(B)** Heatmap of the most DE genes across immune cell subpopulation plotted using cells from the L^nb^ dataset. **(C)** Heatmap of the most DE genes across leukocyte subpopulation plotted using cells from the LCD45^e^ dataset. For all heatmaps, z-scores are shown representing standardized, log normalized expression across cells for a given gene. **(D)** Table of the list of genes used for heatmap analysis for each leukocyte subpopulation. Immune cell subpopulations are grouped from left to right: NK&T, myeloid cells, plasma cells then B cells and the gene lists going from top to bottom are ranked in the same order. The same gene lists were used to analyze cells from each of the three datasets.

### Differential gene expression profiles between datasets reveals pathways affected by tissue processing and establishes that immunologic pathways are preserved

After identification of all the genes that had a statistically significant differential expression (log fold change <0.8 or >1.25 and adjusted *p*<0.05 as shown in volcano plots from Figure 3B) among the three datasets, a canonical pathway analysis was performed using the Ingenuity Pathway Analysis software (IPA, Qiagen, RRID: SCR_008653). The association study was performed for each leukocyte subpopulation: NK & T cells, B cell, plasma cell and myeloid cells. Input of the DE gene name, average log fold change and *p* values provided an output of canonical pathways which were ranked on their likelihood to be altered between pairwise datasets. The probability that a signaling pathway was affected by differences between datasets was represented as a -log(*P* value) and a cutoff of 7.0 was set in order to establish those canonical pathways which were most different. Disease-specific pathways of no relevance to immune-specific subpopulations were excluded (e.g., Atherosclerosis Signaling, Bladder Cancer Signaling) as they were extraneous to this analysis.

Across immune cell subpopulations, there was general agreement between the canonical pathways affected based on the pairwise dataset comparison and therefore NK & T cell subsets and B cell subsets are shown as examples (**Figure 5A-C** and **Figure 5D-F** respectively) as myeloid and plasma cells demonstrated the same pathways which were affected, albeit to varying degrees.

The analysis of DE genes in NK and T cells for LACE^e^ vs L^nb^ pair identified three pathways: EIF2 signaling (-log(*P* value)=14.7, 20 DE genes involved), oxidative phosphorylation (-log(*P* value)=8.8, 11 DE genes involved), and regulation of eIF4 and 70S6K signaling (-log(*P* value)=7, 11 DE genes involved; **Figure 5A**). B cell DE genes between LACE^e^ vs L^nb^, showed a similar result with the addition of mTOR signaling. Of note, many of the genes that led to the presumptive alterations of these pathways were ribosomal proteins with the exception of the oxidative phosphorylation canonical pathway, where genes affected were NADH dehydrogenase subunits, cytochrome c oxidase subunits and ATP synthase subunit F0 subunit 6.

The LCD45^e^ versus LACE^e^ pairwise comparison of canonical pathways was reminiscent of the altered pathways seen in the LACE^e^ versus L^nb^ differential comparison. The expression analysis between L^nb^ and LCD45^e^, which was the pairwise comparison with the best transcriptome concordance and the fewest DE genes, actually had the highest number of altered canonical pathways (**Figure 5C,F**). Even though several canonical pathways were identified as altered between these two datasets (L^nb^ vs LCD45^e^) many of the DE genes that made up the pathway differences were repeated. For instance, ALB or the protein albumin was counted as being a DE molecule in six out of the eight canonical signaling pathways. Likewise, genes such as APOA1, APOA2, ORM1, ORM2 and SERPINA1 contributed to the majority of canonical pathway alterations. In fact, most of the molecules identified were redundant downstream effectors of the pathways and thus do not necessarily reflect true perturbations of the signaling pathway between datasets.

**Figure 5:**
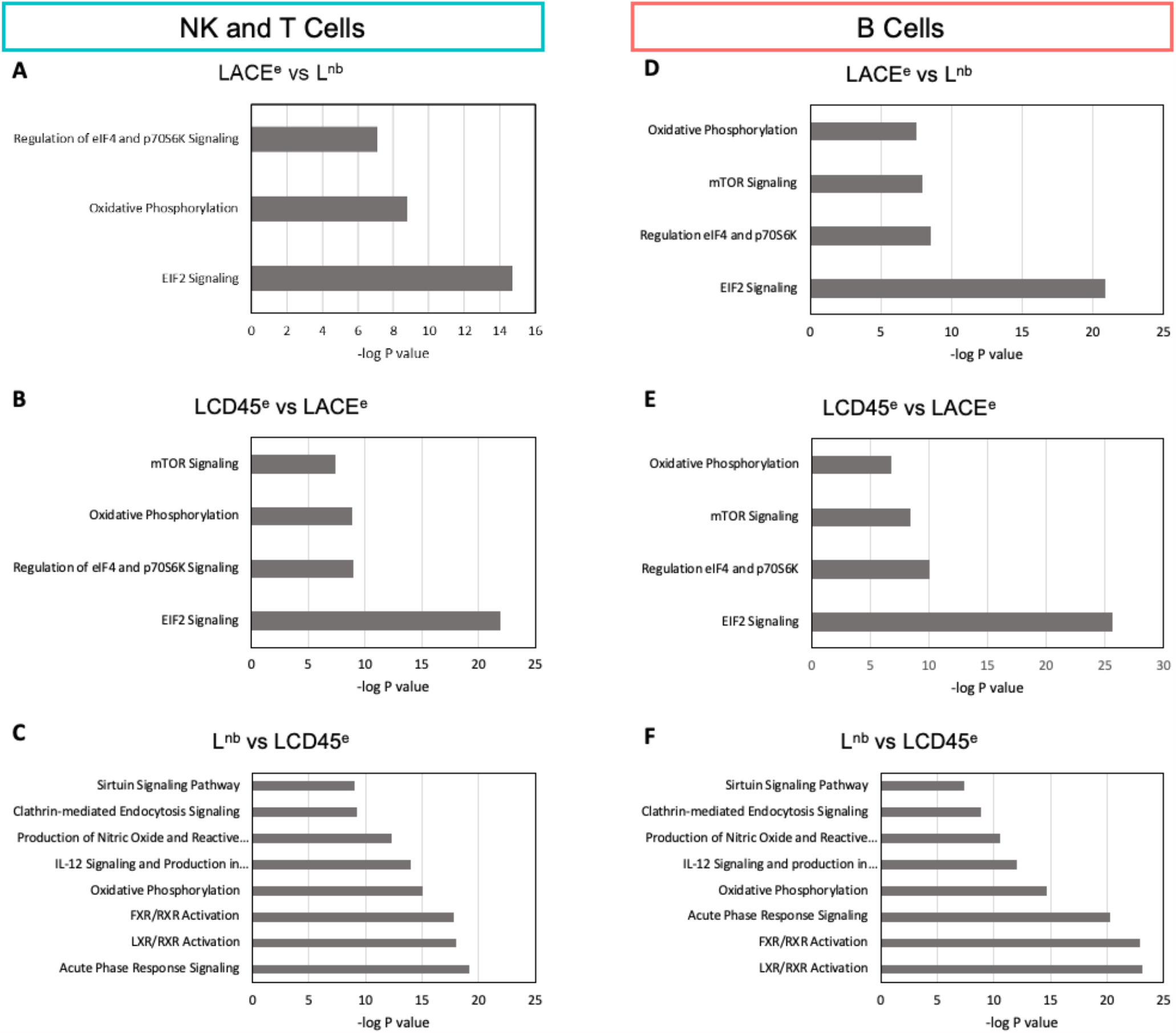
Characterization of differentially expressed genes points to signaling pathways that differ between datasets. **(A**) Pathway analysis of NK and T cells based on differential expression between the LACE^e^ and L^nb^ datasets. **(B)** Identification of canonical pathways altered between LCD45^e^ and LACE^e^. **(C)** Pathway analysis of NK and T cells based on differential expression between L^nb^ and LCD45^e^ in NK and T cells. **(D-F)** B cell pathway analysis as in A-C. The same comparisons were made for myeloid cells and plasma cells and were largely similar to what is shown for the immune subpopulations represented in this figure. Bars represent -log(P values) of each canonical pathway’s likelihood of being altered between comparisons and was obtained with Ingenuity Pathway Analysis Software (Qiagen).

In characterizing the canonical pathways that differed between datasets, the major differences were noted among pathways which involved protein synthesis, cell death and apoptosis signaling. Signaling pathways related to immunologic functions, which are the functions that are of relevance to this study were largely preserved. For example, the T cell receptor signaling pathway, CD28 signaling in T helper cells and IL-6 signaling pathways all had 2 or fewer genes which contributed to differences noted which supports the idea that these pathways were largely comparable between studies and that proceeding with a combined meta-dataset is feasible.

### Part 2: Key features of the integrated liver immune meta-atlas

Given that gene expression correlation was high between datasets and differential gene expression was relatively low, with conservation of phenotypically relevant genes and signaling pathways, we established meta-signatures for each immune cell subpopulation. Genes which were DE from one leukocyte subpopulation versus corresponding cells from a PBMC dataset, were identified. This is in contrast with the prior analysis, which used genes DE between datasets. These candidate genes comprising a list of 449 to 573 total genes depending on leukocyte subpopulation were generated and run through the Ingenuity Pathway Analysis Software (Qiagen, RRID: SCR_008653). The output of this software identified a series of genes with linked cellular functions to determine which canonical pathways were affected.

Pathway analysis output generated characterizations of various cellular biochemical functions. Functions were sub-divided into types including Systemic Autoimmune Syndrome, Signal Transduction, Proliferation, Neoplasia, Inflammation, Infection, Differentiation, Chronic Inflammatory Disorder, Cell Viability, Cell Death, Cell Proliferation, Cell Movement, Binding, and Activation. Z-score values of whether expression was upregulated or downregulated were used to establish a functionality signature for each cell type. Specifically, these pathway analysis data were then transferred for further tertiary analysis in R using the Circlize package to produce meta-signatures as chord plots which relate specific cell functions to individual genes having differential expression. In order to reduce complexity of the figures, redundant cell functions and functions related to disease states noted to be irrelevant to immune subpopulations were excluded. The resulting chord diagrams link genes associated with specific cellular functions, allowing for visualization of the immune phenotype for each leukocyte subpopulation (**Figure 6**).

**Figure 6:**
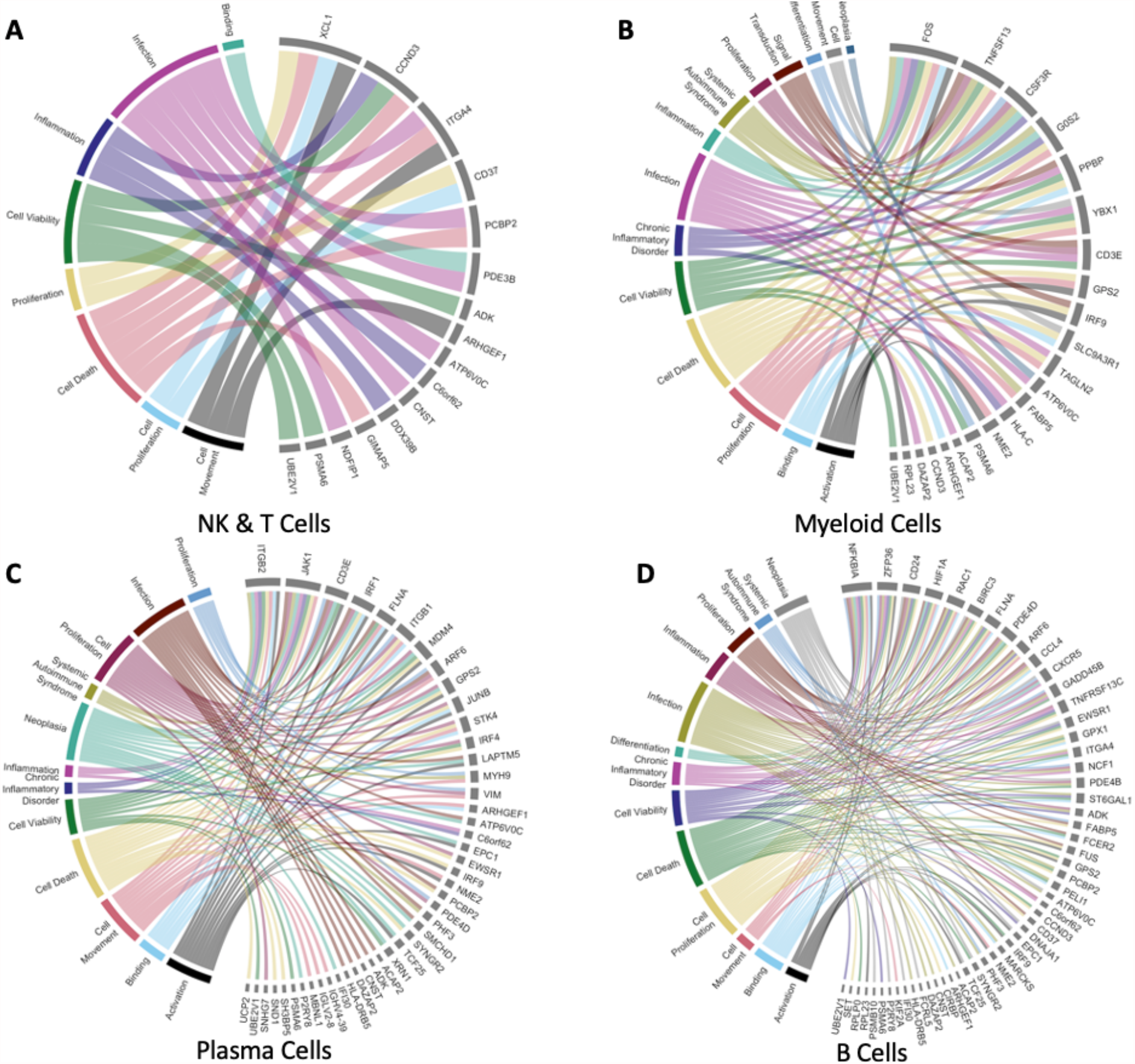
Gene meta-signatures reveal the expression landscape of immune homeostasis in the human liver immune meta-atlas. **(A-D)** Chord plots representing 8-14 immune cell-specific diseases or functions (Systemic Autoimmune Syndrome, Signal Transduction, Proliferation, Neoplasia, Inflammation, Infection, Differentiation, Chronic Inflammatory Disorder, Cell Viability, Cell Death, Cell Proliferation, Cell Movement, Binding, Activation) with links to the respective genes which have been identified as DE between the liver leukocyte subpopulation listed versus the corresponding PBMC leukocyte subpopulation.

NK and T cells represented the largest proportion of cells in our study but had the smallest number of DE genes identified by our analysis, with plasma cells having the most genes. For this immune cell subpopulation, 16 genes were identified as being DE, representing 8 major immunologically relevant diseases and functions (**Figure 6A**). Of the DE genes in NK and T cells, C-C motif chemokine ligand 4 (CCL4L1/CCL4L2), C-C motif chemokine ligand 3 like 3 (CCL3L3) X-C motif chemokine ligand 1 (XCL1), ALOX5AP (involved in leukotriene synthesis) were chemokines and immune modulators noted to be upregulated. Among the most downregulated genes in the NK and T cell subpopulation were LIME1 and TRBV20-1 which are both involved in T cell receptor (TCR) activation signaling. The myeloid lineage was marked by a gene expression signature involving 22 genes across the relevant diseases and functions (**Figure 6B**). Upregulation of the following DE genes was shown: NOP53 ribosome biogenesis factor (involved in apoptosis and cell cycle signaling), Y box binding protein 1 (YBX1, a transcriptional regulator), and TLE5 (transcriptional regulator) among others. The most notably downregulated DE genes were the pro-inflammatory cytokine of the TNF family, TNFSF13, FOS proto-oncogene, CSF3R (GM-CSF receptor controlling proliferation and differentiation of granulocytes; all *p* values < 0.001).

Plasma cells with 45 DE genes shown in the chord diagram demonstrated upregulation of the histone proteins implicated in transcriptional repression (H2BC8 and HDAC3) as well as TLE5, which is implicated in transcriptional regulation and repression (all *p* values < 0.001). The most downregulated genes were immunoglobulin subunit genes from both heavy and light chains: IGHV4-39, IGHV3-11, IGHV3-49, IGHV1-69, IGKV1-16, IGLV2-8, IGLV6-57 (*p* values < 0.001; **Figure 6C**). Lastly, the B cell meta-signature was comprised of 54 DE genes (**Figure 6D**). Upregulation was seen in the chemokine CCL4, SEM1 (part of 26S proteasome complex), NSD3 (transcriptional repression), and BIRC3 (inhibition of apoptosis, all *p* values < 0.001). Genes which were downregulated included immunoglobulin genes: IGKV3-20, IGKV3-15, IGKV1-40, IGHV3-23, IGHV3-30, IGLV1-51, as well as HLA-DRB5 (involved with antigen presentation, all *p* values < 0.001). A detailed report of genes implicated in leukocyte subpopulation homeostasis including log fold change values is summarized in **Supplemental Table 1**.

Identification of the most significantly DE genes pointed to a phenotype of decreased immune cell functioning in human liver immune cell subpopulations versus corresponding peripheral immune cell subpopulations. To explore this further, signaling pathways and functions were characterized using Ingenuity Pathway Analysis (IPA). Histograms were generated for each immune cell subpopulation using the differential expression between PBMC and the human liver meta-atlas in order to characterize which canonical pathways were highly altered between compartments. The significance of the perturbation is charted as -log(P-value) and a cutoff of 2.4 was used in order to identify the most significantly altered pathways. The direction of the perturbation was estimated by a Z-score, which was generated as an output of IPA, and upregulated pathways are color coded as orange, downregulated pathways are blue, and grey pathways correspond to those where the directionality could not be predicted. Heatmaps were also generated which corresponded to specific immune cell functions: Infectious disease, Cell Trafficking, Inflammatory Response, Cell-Cell Interaction, Hematologic development and function, and Cell Death and Survival **(Figure 7)**.

**Figure 7:**
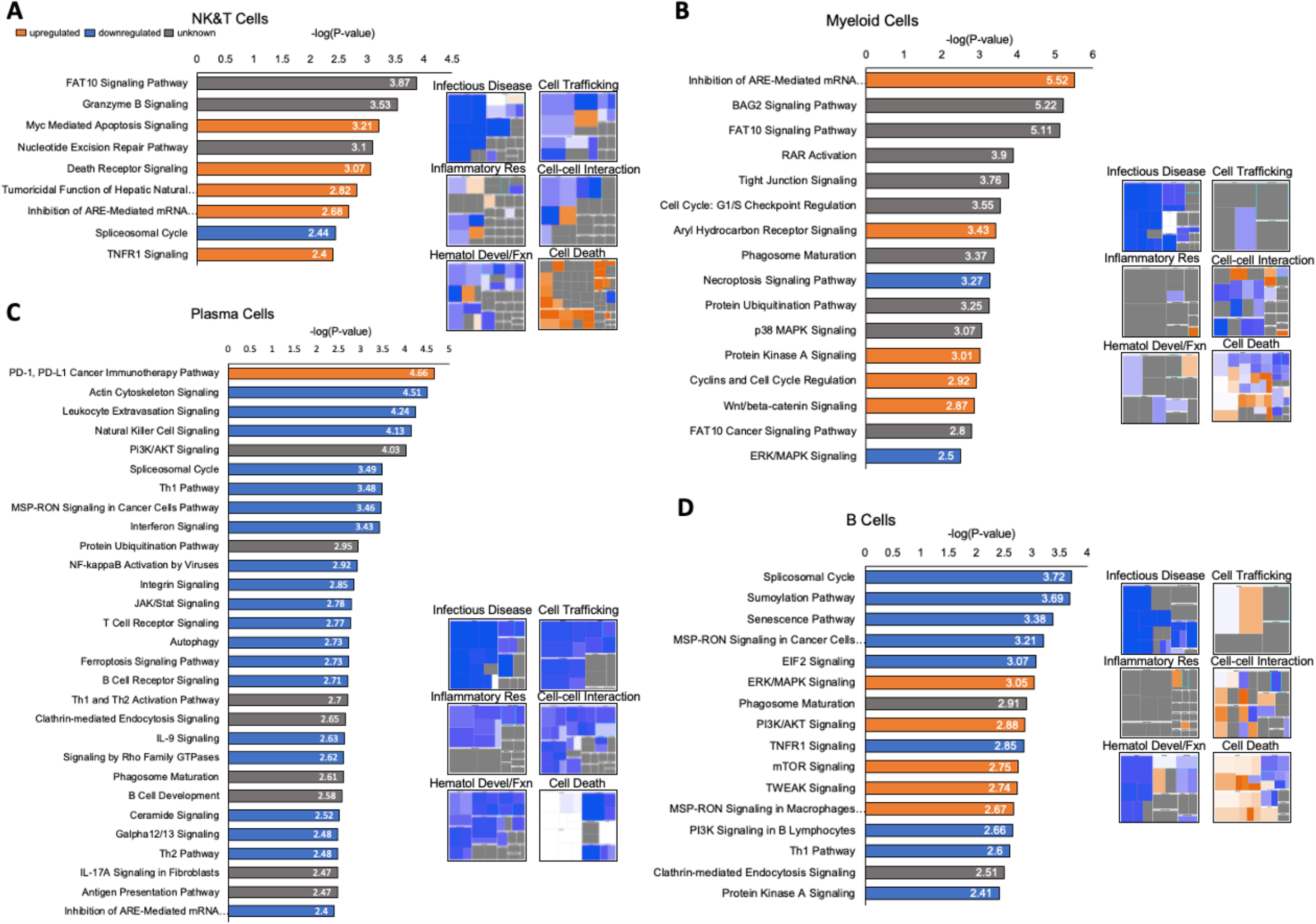
Detailed pathway analysis of human liver immune meta-atlas shows selective upregulation of death-related transcripts and downregulation of immune-related cell functions. **(A)** Pathway analysis of NK and T cells based on differential expression between human liver meta-atlas versus normal human PBMC. Histograms showing -log(P-value) corresponding to the significance of the involved signaling pathway. Orange corresponds to upregulated pathways, blue shows downregulated pathways and gray corresponds to unknown directionality based on Ingenuity Knowledge Base. Heatmaps showing perturbations in specific functions of NK and T cells in the human liver relative to PBMC. **(B)** Myeloid Cell pathway analysis showing canonical pathway functioning in human liver relative to human PBMCs. Heatmaps of specific functions within Myeloid cells relative to PBMCs. **(C)** Canonical pathway functioning Plasma Cells of the human liver relative to PBMC with associated heatmap of specific functions. **(D)** B Cell pathway analysis with histograms showing canonical pathways and heatmaps highlighting specific cellular functions in liver relative to PBMC.

NK and T cells had upregulation of Myc-Mediated Apoptosis Signaling, Death Receptor Signaling, Tumoricidal Function of Hepatic Natural Killer Cells, Inhibition of ARE-Mediated mRNA Degradation Pathway and TNFR1 signaling in the liver compartment relative to PBMCs. Downregulation was demonstrated in the Spliceosomal Cycle. Heatmaps further highlighted immune functions such as infectious disease-related activity, cell trafficking, inflammatory response and cell-cell interactions were largely downregulated, whereas cell death and apoptosis functions were upregulated **(Figure 7A)**. In the myeloid cell subpopulation, upregulated canonical pathways were Inhibition of ARE-Mediated mRNA Degradation Pathway, Aryl Hydrocarbon Receptor Signaling, Protein Kinase A Signaling, Cyclins and Cell Cycle Regulation, and Wnt/beta-catenin Signaling. Downregulated pathways were Necroptosis Signaling and ERK/MAPK Signaling. Phenotypic patterns of diminished immune cell functioning, and enhanced cell death were seen in this population as well **(Figure 7B)**.

Plasma cells had upregulation of the PD-1/PD-L1 Cancer Immunotherapy Pathway and downregulation of a substantial number of canonical pathways. Heatmaps showed that phenotypically these cells were largely in a state of depressed functioning relative to the peripheral blood immune compartment (Figure 7C). Human liver B cells, comprising the smallest immune cell subpopulation, showed upregulation of ERK/MAPK Signaling, PI3K/AKT Signaling, mTOR Signaling, TWEAK Signaling and MSP-RON Signaling in Macrophages Pathway. Downregulation was demonstrated in the Spliceosomal Cycle, Sumoylation Pathway, Senescence Pathway, MSP-RON Signaling in Cancer Cells Pathway, EIF2 Signaling, P13K Signaling in B Lymphocytes, Th1 Pathway and Protein Kinase A Signaling. Heatmap analysis showed a mixture of upregulated and downregulated functions. Infectious disease and hematologic development related functions were again depressed, but cell trafficking and inflammatory responses had a modest upregulation in liver relative to PBMC. Cell death functions were again enhanced. Overall, these results indicated that liver immune homeostasis points to a general phenotype of depressed immune and pro-inflammatory functions via the aforementioned pathways and a possible increase in cell death and apoptotic events.

## DISCUSSION

In this study, meta-analysis of three high-quality scRNA seq studies enabled deep characterization of liver immune cell function and features related to physiologic immunotolerance and immune homeostasis. Focused examination of immune cell types, proportions, and gene expression profiles of revealed that significant differences exist between datasets created from ‘normal human liver’ samples. Despite these differences, the overall ranking of abundance of immune subsets was preserved, with NK and T cells dominating and plasma and B cells being relatively rare. Further analysis of gene expression profiles involving the most DE genes in the integrated dataset provided the opportunity to minimize the effect of technical ‘noise’ and establish the gene expression signature of human liver immune milieu. These results point to reduced expression of immune related functions and enhanced expression of cell death-related pathways. This human liver meta-atlas has the potential to serve as a key reference point for future studies using scRNA seq designed to characterize immune-based pathologies within the liver, such as liver allograft rejection, autoimmune disease, and the tumor microenvironment in primary liver malignancies.

To establish validity of performing a meta-analysis of an integrated dataset using scRNA seq data generated in three distinct labs with different tissue handling, processing, and sequencing techniques, visualization of the integrated data with co-clustering of cell populations was performed (**Figure 2**). This approach was based on the work of Bonnycastle et al, who specifically studied the impact of tissue processing and handling on the integrity of scRNASeq data.(16) In our analysis, immune cell proportions and expression profiles differed somewhat between datasets, with highest correlation between the L^nb^ and LCD45^e^ studies (**Supplemental Figure 5**). This is not particularly surprising, as these two datasets used the same 10x Genomics technique for the scRNA seq. It is important to note that extraction of the CD45+ population and combination with other non-liver derived CD45+ cells did not appear to impact the integrity of the data, as was the case with the LCD45^e^. While various scRNA seq techniques have the potential to introduce noise and bias into the transcriptome data, the Seurat pipeline has been designed to correct for some of these differences by re-aligning cell clusters based on nearest neighbor correction and allowing for integrated downstream analyses.(18,19) This would suggest that differences we observed in expression analyses could be the result of pre-analytical variables such as tissue processing. We further characterize noise with the quantification of an established set of human housekeeping genes.(28) There were differences noted in expression of human housekeeping genes within an immune cell subpopulation between datasets, however, mean expression values were not substantially altered and differences noted here are likely the result of large sample sizes. A potential explanation for differences in cellular proportions may be related to sample size effect as well. The LACE^e^ dataset encompasses the largest sample of human tissues, derived from nine total liver specimens, which likely introduces a higher degree of biological variation than L^nb^ and LCD45^e^ which have five and three liver donors, respectively. The clinical scenario also differs between LACE^e^, which was created using partial hepatectomy specimens from patients with liver malignancy, and the other two studies, which involve specimens derived from adult brain-dead organ donors. Despite attempts at correction, there were differences noted in gene expression analysis, yet immune subpopulations still co-clustered with good interdigitation between datasets and DE analysis showed good agreement with heatmaps (**Figures 2 and 4**). As such, we proceeded with deeper immune profiling using the integrated dataset.

Differences in tissue handling and processing has been shown to prompt transcriptional changes within the cell.(15,16,31) A major novelty of our study is the in-depth characterization of differential expression between datasets with identification of relevant canonical pathways affected. Our results indicate that pathways involved in oxidative phosphorylation and cell stress were primarily affected and that immunologically relevant pathways were largely preserved. We suspect that non-immune function related changes may be the consequence of pre-analytical tissue processing. Known pre-analytical variables related to each technique used for the generation of each dataset are summarized in **Table 1**. Unknown variables of potential importance could include the time interval of ‘Pringle’ clamp time during resection or from the moment of tissue resection to cell processing and sequencing, which could lead to significant ischemic insults and thus impact transcription profiles. In ischemic injury of rodent and human cortex, insult has been demonstrated to promote pro-inflammatory gene expression alterations.(32,33) The fact that these ischemia-induced pro-inflammatory changes are not demonstrated in this study shows that either ‘Pringle’ time in liver specimens captured in these datasets were either not significant enough to cause ischemia, or that liver ischemia does not produce as profound an inflammatory response as cortical ischemia. Overall, the observation that immune pathway profiles for each subpopulation were relatively conserved supports the use of this novel meta-analysis approach to examine existing datasets in order to understand immune homeostasis in human livers.

Immunotolerance is a unique feature of hepatic homeostasis and is thought to be a consequence of the constant stream of antigens being delivered to the liver from the gut. This involves regulation of both the adaptive and innate immune systems to prevent an uncontrolled inflammatory cascade in response to foreign antigens in the portal circulation.(1,34) One potential theory is that effector T-cells are regularly destroyed in the liver. Animal studies have suggested that hepatic immune regulation is mediated by the milieu of anti-inflammatory cytokines and inhibitory regulators such as IL-10 and TGF-ß, leading to inhibition of effector T cell function and expansion of the Treg population.(35) Comparison of hepatic immune cell gene expression patterns to a different immune compartment enabled identification of a liver-specific signature of immune homeostasis. Indeed, this analysis revealed that NK, T cell, and myeloid subpopulations had enhanced cell death functions and depressed immune-related functioning (Figure 7). Similarly, plasma and B cells both showed downregulation of immunoglobulin genes and exhibited downregulation of immune functional pathways.

There are limitations to this study. Further differences in scRNA seq technique (comparing 10x Genomics to mCel-seq) likely introduces a differing amount of bias into the sequencing datasets. In addition, because this study explored pre-existing datasets, there was no ability to control for patient-specific parameters in this study design such as inclusion/exclusion based on liver function studies or other potentially relevant clinical factors including duration and storage of samples prior to processing. Despite this, combining scRNA seq and other genomics datasets for the characterization of normal physiology and disease processes will become a more prevalent practice and further efforts should be made to standardize the processes of pre-analytical processing variables as well as to enhance the integration abilities during data analysis.

In conclusion, our results present an integrated dataset of scRNA seq results of the liver immune environment from ∼32,000 cells across 17 human livers, making it the largest human liver meta-atlas available. Key immune subset expression profiles that describe the landscape of liver immune homeostasis were defined and can be explored in real-time on our interactive website.(27) These results can be incorporated into future RNA sequencing studies and have implications for understanding mechanism of disease and identifying new therapeutic targets in immunologic diseases of the liver including allograft rejection and immune tolerance.

## METHODS

Institutional Review Board approval was not required for this study, as deidentified, publicly available data obtained from human subjects was analyzed. Data is available in the public domain as outlined below.

### Systematic review and data acquisition

A comprehensive review of the literature for scRNA seq studies involving normal human liver yielded three recent publications: 1. Aizarani *et. al*., and based on the methods used in the study, it was referred to as “Liver Atlas, Cholangiocyte and Endothelial enriched” (LACE^e^), 2. Zhao *et. al*., referred to as “Liver CD45+ enriched” (LCD45^e^), and 3. MacParland *et. al*., referred to as “Non-biased Liver” (L^nb^).(7–9) Each raw dataset was found in the Gene Expression Omnibus, LACE^e^: GSE124395, LCD45^e^: GSE125188, L^nb^: GSE115469.(17) Datasets were imported into R as UMI count matrices using the RStudio interface. The workflow for isolation of our cell populations of interest from each dataset is summarized in **Figure 1**. Briefly, the CD45+ cell compartment was isolated in the LACE^e^ and L^nb^ studies in order to extract the leukocyte population and exclude hepatocytes and other non-parenchymal cells. For LCD45^e^, the CD45 population was isolated prior to sequencing, and included cells derived from liver, spleen, and peripheral blood that had been barcoded for source identification. In this case, only the liver cells were extracted for analysis. Authors from each dataset were contacted for clarification on aspects of the datasets as needed in order to ensure the accuracy of our analysis.

### Data analysis and visualization

The Seurat version 3.0 R toolkit (RRID: SCR_007322) was used for all data analysis due to its ability to integrate across datasets; the data analysis pipeline followed the tutorial outlined by the package developers.(18,19) Single cell expression count matrices barcodes and gene IDs from each study were downloaded from GSE. After import of the datasets into Seurat, we filtered barcodes for CD45+ when handling the LACE^e^ and L^nb^ datasets. The LCD45^e^ dataset included cell counts which were exclusively from CD45+ cells, therefore this filtering was not necessary. Spleen and peripheral blood mononuclear cells (PBMCs) were handled by excluding the associated barcodes and only including barcodes for the liver immune cells. To normalize the data, UMI counts were scaled by library size and a natural log transformation; gene counts for each cell are divided by the total UMI count of that cell, scaled by a factor of 10,000, and then transformed via a natural log plus 1 function (“NormalizeData”). For downstream analysis, normalized data is additionally scaled so that the mean expression across cells is 0 and the variance is 1 (“ScaleData”). In order to reduce the dimensionality of the data for clustering functions, Principal Component Analysis (PCA) was utilized, and we determined the first 30 principal components explained sufficient observed variance (“RunPCA”). Next, to identify clusters within the reduced dimensional space, cells were embedded in a k-nearest neighborhood-based graph structure (“FindNeighbors”) and were then partitioned into clusters (“FindClusters”). Finally, to aid in visualization, Uniform Manifold Approximation and Projection (UMAP) was run over the reduced dimensional space (‘RunUMAP”) and identified clusters were projected on to the UMAP plot. Accounting for noise and batch differences between the three studies was done by using reference cells from each. These anchors were identified (“FindIntegrationAnchors”), which represent pairs of cells from each dataset identified and scored based on their close proximity using k-nearest neighborhood approach. These anchors are then used to measure the expression difference between studies (“IntegrateData”), which is then removed from the corresponding normalized data. Integration is run between the normalization and scaling steps.(19)

Clustering was performed in Seurat v3. To establish reliability of this method, the clustering algorithm was compared between Seurat and an alternative approach using the Harmony R package (https://github.com/immunogenomics/harmony). This package scales the data to make nearby cells more similar when clustering is performed. Harmony is also easily incorporated into the Seurat pipeline. Results using Harmony-based clustering were essentially identical to Seurat. Across cell types, 96.5% to 99.9% of cells in Seurat clusters were present in the same Harmony clusters (**Supplemental Figure 1**). Additionally, agreement was high across datasets ranging from 96.4% to 99.4%. This data confirms that our clusters are accurate and downstream analyses should not be affected by the choice in algorithm.

Utilizing the UMAP plot of the integrated data, we were able to manually identify four clusters corresponding to four major immune subpopulations by plotting expression of known biomarkers and grouping previously identified clusters: myeloid cells, NK&T cells, B cells, and plasma cells (**Supplemental Figure 2**). CD3D was used to identify T lymphocytes, and NK cells were identified by expression of KLRF1 and FCGR3A. The myeloid cell lineage was identified using FCGR3A and the specific marker, CD14. The B cell cluster was classified using the CD19 marker, and plasma cells were identified by SDC1 (CD138) expression.(21–25) After clustering, immune cell proportions were quantified and characterized between datasets using a Chi-squared test (*α*=0.05). Gene correlation analysis was performed between datasets as pooled cell types and additionally with stratification by major cell type using both standard linear regression and rank-rank hypergeometric overlap (RRHO).

### Differential expression analysis

DE genes were identified across immune cell subpopulations (i.e., between a particular cell subpopulation and all other cell subpopulations) within a dataset. Log normalized gene expression were used, averaging over all cells in a given subpopulation for a specific gene and taking the difference to the same of all other cell subpopulations (“FindMarkers”), using the Wilcoxon rank-sum test for significance. Genes that were DE were examined both within an immune cell subpopulation and between datasets. Similarly, DE genes were identified across datasets (i.e., pairwise between datasets) within a particular cell subpopulation using the same methods. To account for some technical variation between dataset, we calculated the difference of differences of a particular subpopulation between datasets and all other cell subpopulations between datasets. Log normalized gene expression was used as described above taking the difference of differences between dataset (pairwise) and cell subpopulation (particular versus all other), using a t-test for significance.

The candidates for immune profile of human liver across our major cell types of interest were identified by applying differential expression analysis across immune cell subpopulations in liver in the combined meta-atlas compared with immune cell subpopulations from a published dataset generated from normal human PBMC (67,221 cells; NCBI Gene Expression Omnibus: GSE171555).

### Pathway analysis

Gene IDs, expression fold changes and p-values generated from differential expression analyses were imported into Ingenuity Pathway Analysis software (IPA, Qiagen, RRID: SCR_008653). Core expression analysis was performed which generated canonical pathways which were upregulated or downregulated based on these data. Gene function heatmaps were also generated in order to provide a comprehensive expression profile.

### Volcano plots

Pairwise comparisons of genes between data sets or cell types were conducted using Seurat “FindMarkers” function with default Wilcoxon Rank Sum test. Volcano plots were constructed using these results; genes were identified and colored based on P-adj_Bonfcorr_<0.05 and a ratio of expression (exponentiated log fold change) ≥1.25 or ≤0.8. Additionally, differences of differences were calculated in a similar way comparing the differential expression between datasets and the differential expression between immune cell subpopulations. T-tests were performed to determine significance and genes were colored based on the previous criteria plus P-adj_Bonfcorr_<0.05 of the difference of differences effect.

### Correlation analysis

Expression levels were averaged over cells and plotted pairwise between datasets. Scatter plots were fit with linear and quadratic regression models to show potential relationships. Additionally, genes were ranked based on their differential expression between cell types and these rankings were compared between datasets. Correlation between the rankings was assessed multiple ways. First via scatter plots and Spearman’s correlation coefficient. Second using Rank-Rank Hypergeometric Overlap from the RRHO package in R and constructing heatmaps showing regions of significant overlap between the ranked lists.(26)

### Heatmaps

The top DE genes between cell types were identified and combined between each dataset. Raw expression values for these genes were then normalized by subtracting the mean expression and dividing by the standard deviation of a given gene over all cells. Heatmaps were constructed based on these values; identified genes are shown as rows while cells are shown as columns. Cells are grouped based on cell type while genes are grouped based on the cell type differential expression from which they were identified. Outputs from heatmap analyses were used in the creation of an online interactive tool of the combined human liver meta-atlas.(27)

### Chord plots

Results from the pathway analysis were used to select cell functions that were relevant to the immune subpopulations. Cell functions and genes were linked to represent the presence of a connection between the two, yes/no. From there, plots were constructed (Circlize “chordDiagram”) by connecting selected cell functions to the genes that were identified performing them, grouping by cell function.

## DATA AVAILABILITY

The datasets analyzed for this study can be found in the Gene Expression Omnibus, LACE^e^: GSE124395, LCD45^e^: GSE125188, L^nb^: GSE115469, PBMC: GSE171555 (normal samples GSM5227141, GSM5227117, GSM5227130, GSM5227148). An online “point-and-click” resource for interacting with the meta-atlas which is available on our lab website: https://usctransplantlab.shinyapps.io/meta_rna_seq/

## CODE AVAILABILITY

R code used for data analysis and creation of graphics is available upon request and can be accessed via github.

## ACKNOWLEDGEMENTS

We thank the Eli and Edythe Broad Center for Regenerative Medicine at the University of Southern California, Keck School of Medicine for providing the Broad Clinical Research Fellowship which was awarded to Dr. Rocque for 2020-2021. We also thank the USC Libraries Bioinformatics core for providing software licensing and resources.

## AUTHOR INFORMATION

BR contributed to study design, acquiring data, analyzing data, and writing the manuscript. AB contributed to study design and writing the manuscript. PS was responsible for acquiring initial datasets and performed computational analyses and data integration. CG contributed to study design, analyzing data, statistical analyses and aided with writing the manuscript. DGH helped with study design and writing the manuscript. YEL provided resources for data analysis and contributed to study design. NU provided insights for study design and contributed to verification of data analysis techniques. JL and OA helped with the initial study design. JE conceived the original idea, supervised the project and contributed to study design and data analysis.

