## Supplemental Data for "Creation of a single cell RNA seq meta-atlas to define human liver immune homeostasis"

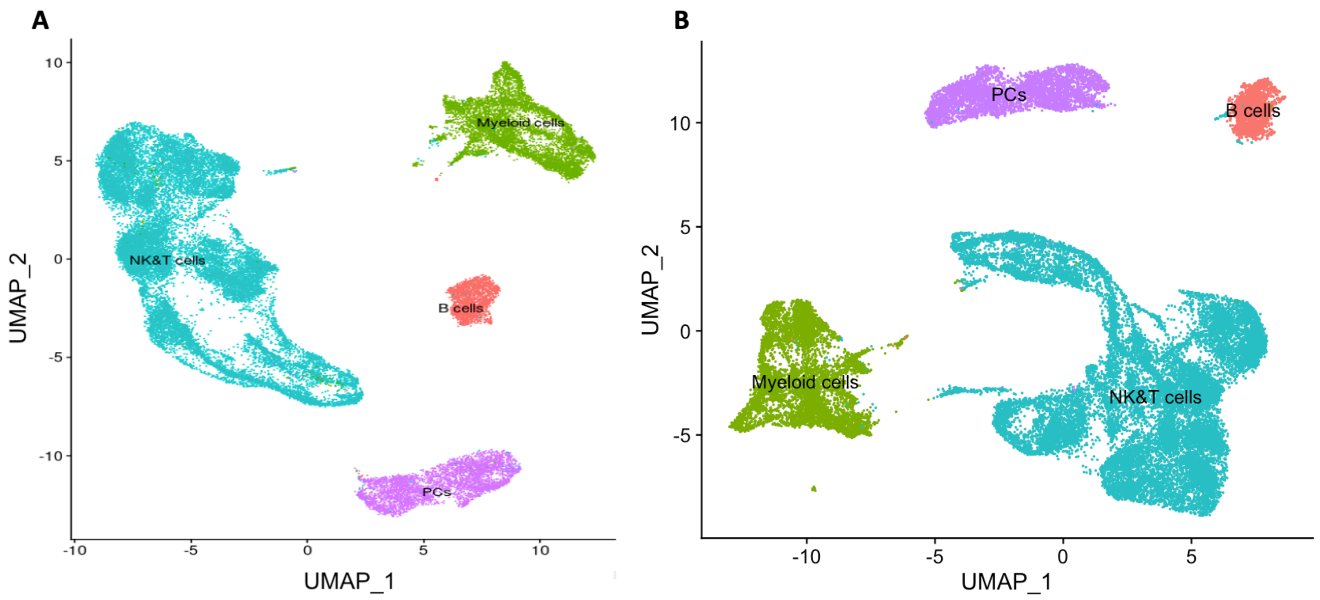


**Supplemental Figure 1: Comparison of clustering algorithms for identification of immune cell subpopulations.**

**(A)** Clustering of 4 major immune cell subpopulations among the three datasets of interest using the Seurat v3 clustering algorithm. **(B)** Clustering of the same datasets using the Harmony algorithm. Comparison across cell subpopulations revealed 96.5% to 99.9% agreement between Seurat and Harmony. Agreement across datasets was 96.4% to 99.4%.

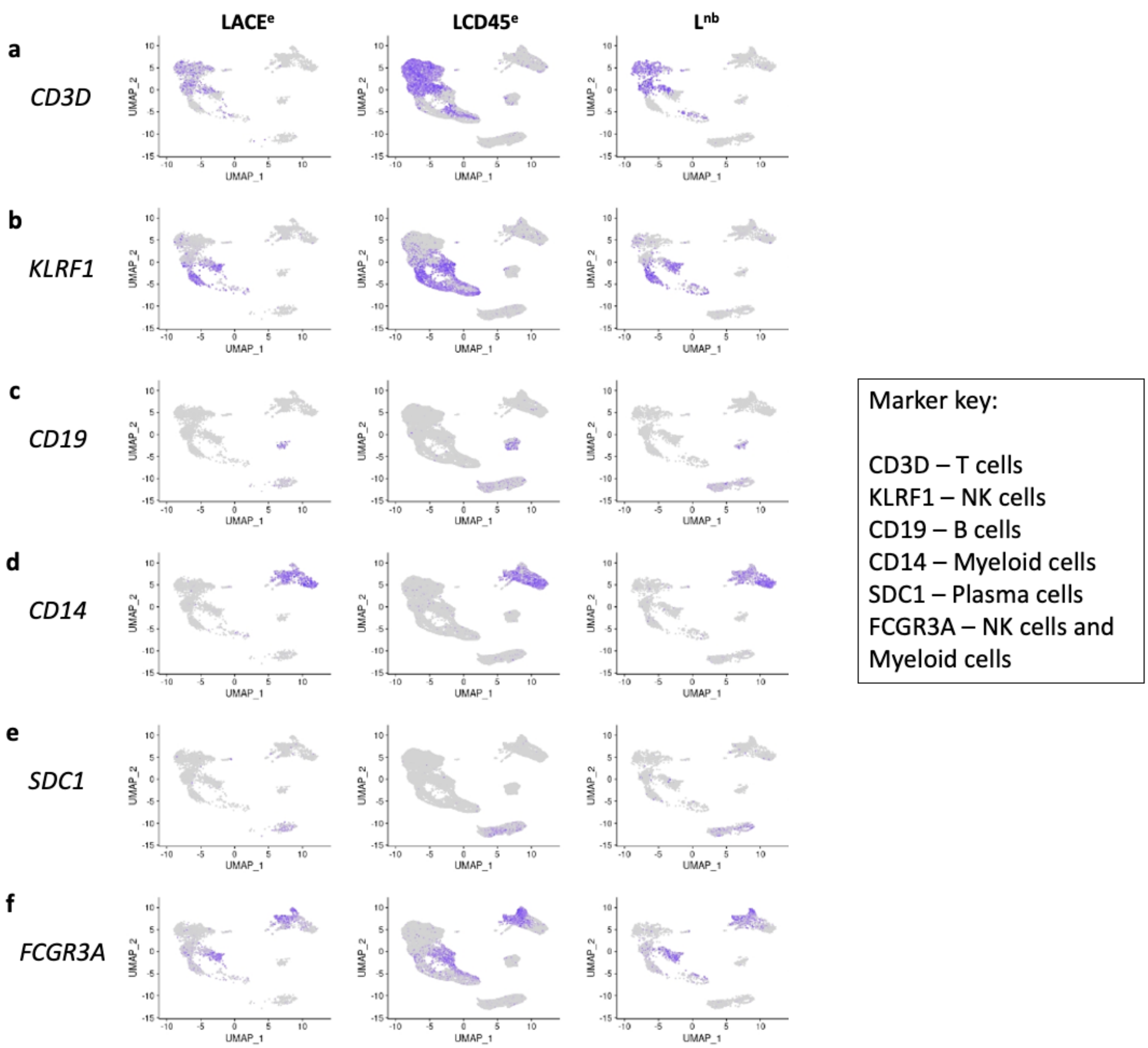


**Supplemental Figure 2: Identification of cell clusters using cell marker feature analysis.** **(A)** CD3D-expressing cells are shown as purple dots and specifically represent T lymphocytes. **(B)** KLRF1 is used to distinguish the clustering of NK cells. **(C)** CD19, a known marker for B lymphocytes, identifies the unique clustering of this cell population. **(D)** CD14-expressing cells identify the cluster with cells from the myeloid lineage. **(E)** SDC1, used to identify the plasma cell cluster. **(F)** FCGR3A expression is seen in both NK cells and myeloid cells.

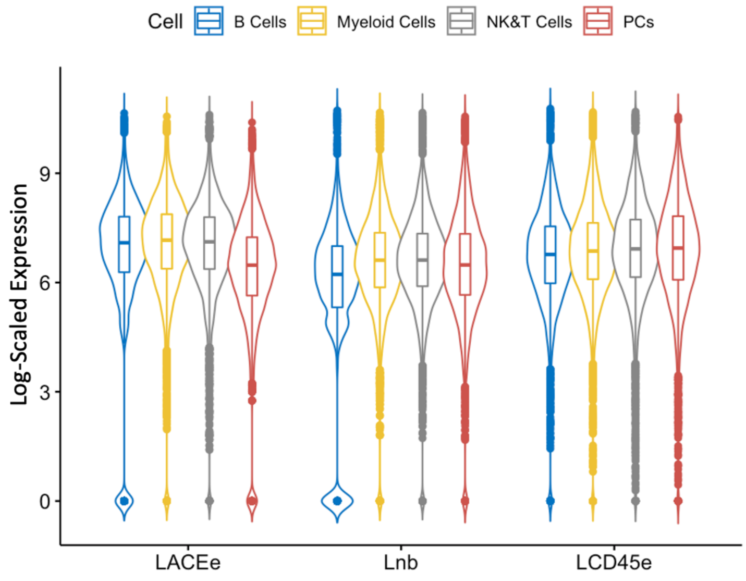


***Table of Means and Standard Deviation***

| | LACEe | Lnb | LCD45e |
|---|---|---|---|
| B Cells | 6.73 (1.961) | 5.71 (2.289) | 6.69 (1.452) |
| Myeloid Cells | 7.01 (1.426) | 6.57 (1.399) | 6.82 (1.41) |
| NK&T Cells | 6.97 (1.384) | 6.56 (1.379) | 6.86 (1.436) |
| PCs | 6.27 (1.676) | 6.43 (1.552) | 6.87 (1.498) |

***Pairwise Mean Comparisons***

| Cell Type | Source 1 | Source 2 | Adjusted P-value |
|---|---|---|---|
| B Cells | LACEe | LCD45e | 0.123 |
| B Cells | LACEe | Lnb | <.01 |
| B Cells | LCD45e | Lnb | <.01 |
| Myeloid Cells | LACEe | LCD45e | <.01 |
| Myeloid Cells | LACEe | Lnb | <.01 |
| Myeloid Cells | LCD45e | Lnb | <.01 |
| NK&T Cells | LACEe | LCD45e | <.01 |
| NK&T Cells | LACEe | Lnb | <.01 |
| NK&T Cells | LCD45e | Lnb | <.01 |
| PCs | LACEe | LCD45e | <.01 |
| PCs | LACEe | Lnb | <.01 |
| PCs | LCD45e | Lnb | <.01 |

**Supplemental Figure 3: Comparison of expression levels among housekeeping genes.** An array of log-scaled expression values for a pre-determined set of housekeeping genes were characterized in each cell type and in each individual dataset. These are represented as violin plots showing the density curve with boxplots showing the median, 25$^{th}$ and 75$^{th}$ percentiles, and whiskers for 1.5*IQR, and are color coded according to cell type. Mean expression levels of all housekeeping genes are shown in the “Table of Means and Standard Deviation”. These reveal relatively similar expression levels although further analysis did show significant differences between pairwise comparisons (one-way ANOVA test).

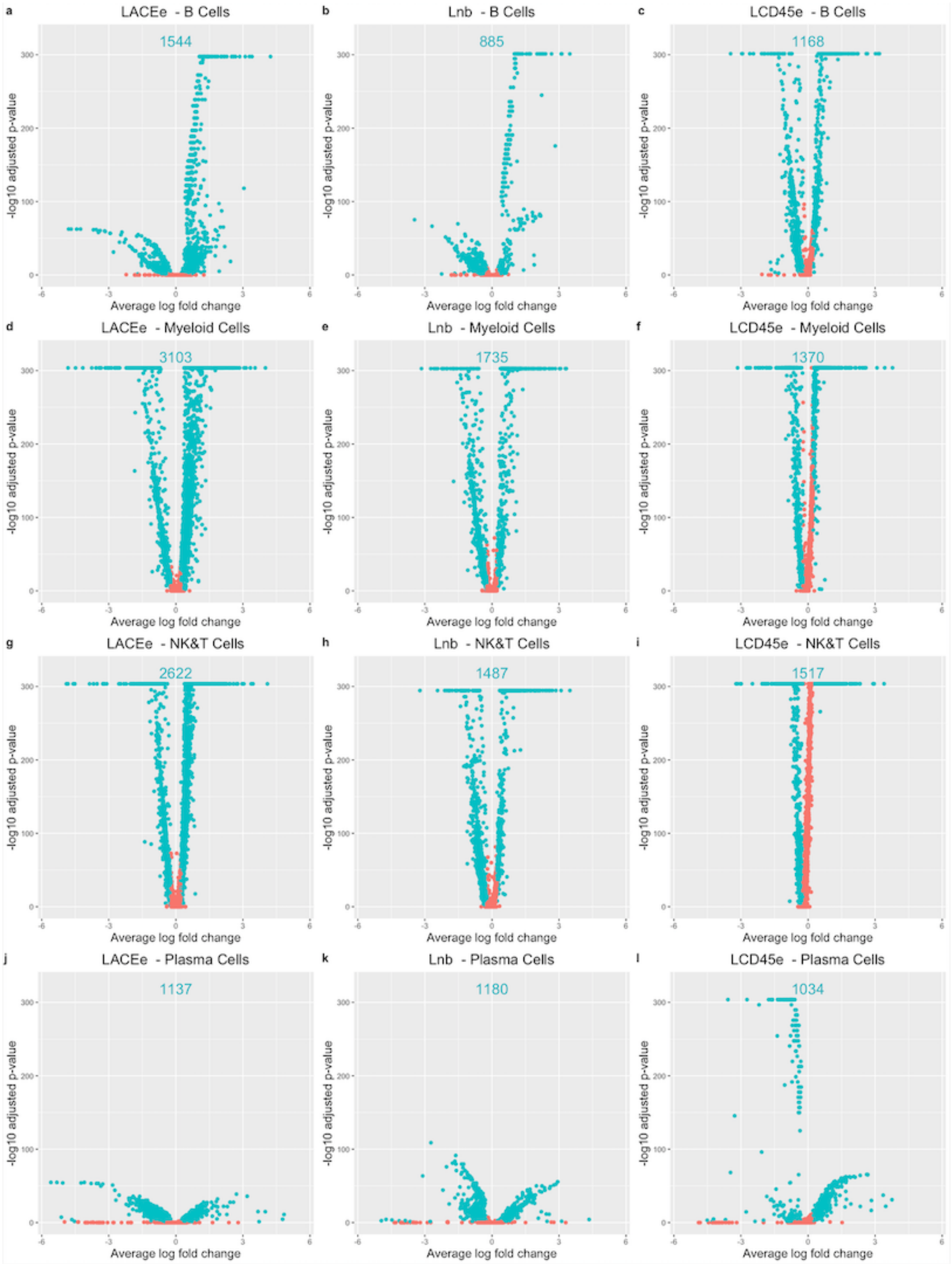


**Supplemental Figure 4: Quantification of the number of differentially expressed genes between immune cell subpopulations within datasets.**

**(A-C)** Volcano plots of DE genes between B cells and all other cell types for each one of the normal human liver scRNA seq studies ($\text{LACE}^{e}$, $\text{L}^{nb}$ and $\text{LCD45}^{e}$ respectively). DE genes are represented in blue and were noted to have a fold-change ratio of less than 0.8 or greater than 1.25. Non-DE genes are plotted in red. **(D-E)**. Volcano plots of DE genes when comparing cells of the myeloid lineage to all other cell types. **(G-I)** Volcano plots of DE genes in NK&T cells versus all other cell types. **(J-L)** Volcano plots of DE genes in plasma cells versus all other cell types.

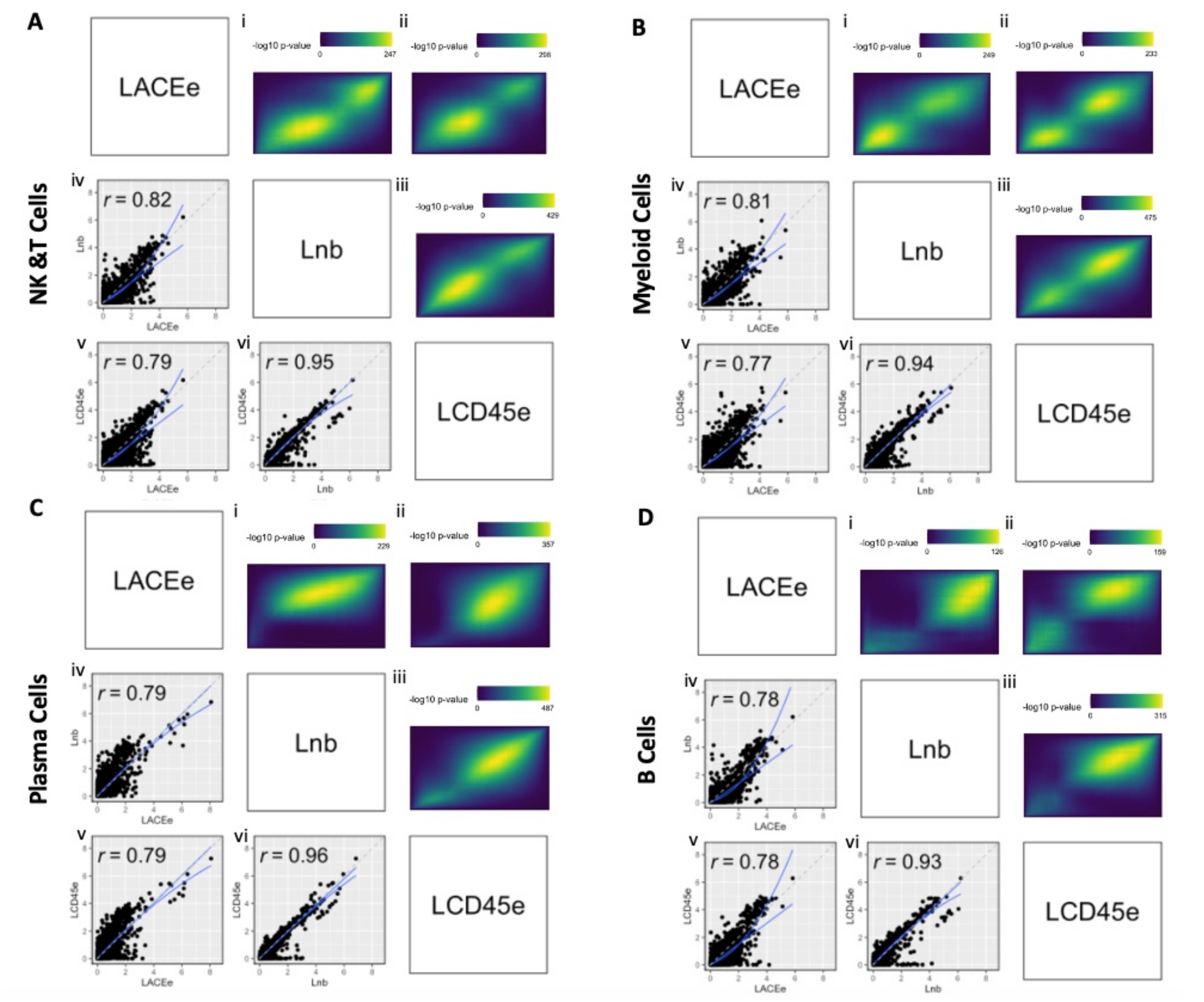


**Supplemental Figure 5: Correlation of gene expression based on immune cell subpopulation.**

Pairwise expression correlation analysis was made for each cell type. **(A)** Gene expression among NK and T cells was compared between datasets. The black and gray panels show how well genes correlate between the three datasets and the dashed diagonal line represents an idealized relationship. The straight solid blue line represents the linear best fit and the curved solid blue line shows the quadratic relationship with linear correlation coefficients listed. Colored diagrams show heatmaps of the rank-rank hypergeometric overlap. Yellow regions indicate that for these pairwise comparisons, NK and T cells have a higher amount of agreement or overlap among the more highly expressed genes (lower left quadrant of the graph) and also with a good amount of agreement in genes that are expressed at lower levels (top right quadrant of the graph). **(B)** The same analysis is repeated for myeloid cells. RRHO shows differing amount of overlap depending on the ranked gene expression. **(C)** Correlation analysis for plasma cells. The best correlation is among genes with mid-to-low ranked expression. **(D)** Correlation analysis for B cells. Correlation is maximized in the genes expressed at lower levels.

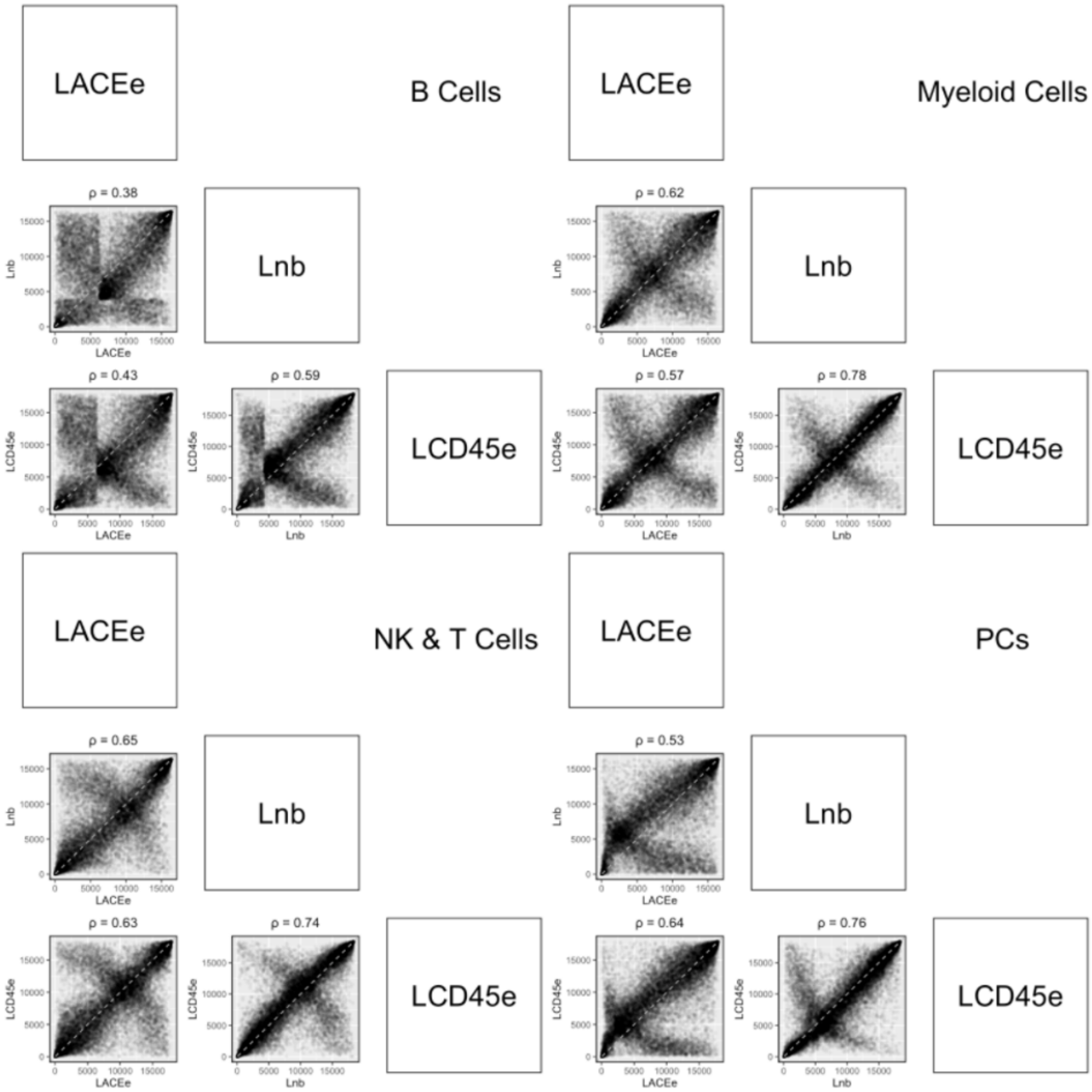


**Supplemental Figure 6: Rank-Rank Hypergeometric Overlap analysis.**
RRHO analysis showing correlation plots between studies in B cells, myeloid cells, NK & T cells and plasma cells.

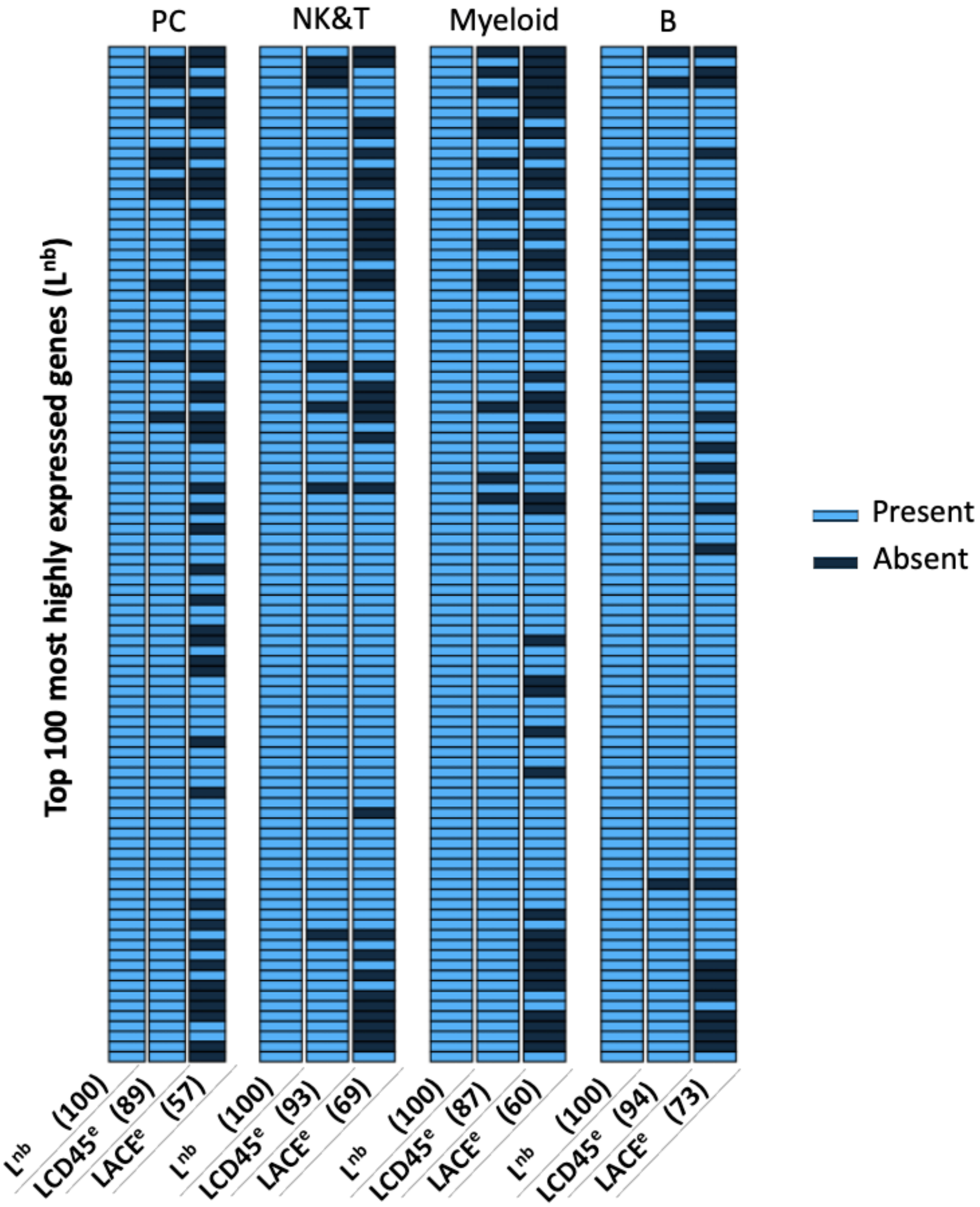


**Supplemental Figure 7: Comparison of the 100 most abundant genes expressed, stratified by immune cell subpopulation and compared across datasets.**

Top 100 most abundant genes were identified for each major cell group in the $L^{nb}$ paper. The presence of these same genes among the top 100 list for $LCD45^{e}$ and $LACE^{e}$ are noted by a light blue box. If the gene was not among the top 100 most abundantly expressed genes in the dataset listed, it is denoted by a dark blue box.

**Supplemental Table 1: Gene expression signatures associated with each leukocyte subpopulation**

| Subpopulation | Gene symbol | Up/Down-regulation | Log_FC | P value |
|---|---|---|---|---|
| **NK & T Cells** | | | | |
| | ATP5F1 | Upregulated | 1.004 | <0.001 |
| | ATP6V0C | Downregulated | -1.305 | <0.001 |
| | CCL3L3 | Upregulated | 0.882 | <0.001 |
| | CCL4L2 | Upregulated | 1.107 | <0.001 |
| | DDX39B | Downregulated | -0.882 | <0.001 |
| | GLTSCR2 | Upregulated | 1.866 | <0.001 |
| | LIME1 | Downregulated | -1.4 | <0.001 |
| | NME2 | Downregulated | -1.343 | <0.001 |
| | P2RY8 | Downregulated | -1.2 | <0.001 |
| | PCBP2 | Downregulated | -1.006 | <0.001 |
| | PDE3B | Downregulated | -1.038 | <0.001 |
| | RPS20 | Upregulated | 1.91 | <0.001 |
| | SELT | Upregulated | 1.057 | <0.001 |
| | SEPT1 | Upregulated | 0.993 | <0.001 |
| | TRBV20-1 | Downregulated | -1.103 | <0.001 |
| **Myeloid Cells** | | | | |
| | ATP5H | Upregulated | 1.105 | <0.001 |
| | ATP6V0C | Downregulated | -2.145 | <0.001 |
| | FAM26F | Upregulated | 1.296 | <0.001 |
| | G0S2 | Downregulated | -1.843 | <0.001 |
| | GLTSCR2 | Upregulated | 1.238 | <0.001 |
| | LINC00936 | Upregulated | 0.947 | <0.001 |
| | NME2 | Downregulated | -1.784 | <0.001 |
| | RPL23 | Upregulated | 1.343 | <0.001 |
| | RPS20 | Upregulated | 1.862 | <0.001 |
| | TRAPPC5 | Downregulated | -1.437 | <0.001 |
| **Plasma Cells** | | | | |
| | AC233755.2 | Downregulated | -2.767 | <0.001 |
| | ATP6V0C | Downregulated | -1.045 | <0.001 |
| | GLTSCR2 | Upregulated | 1.008 | <0.001 |
| | IFI30 | Downregulated | -1.088 | <0.001 |
| | IGHV1-69 | Downregulated | -1.989 | <0.001 |
| | IGHV3-11 | Downregulated | -4.506 | <0.001 |
| | IGHV3-49 | Downregulated | -3.679 | <0.001 |
| | IGHV3-7 | Downregulated | -4.445 | <0.001 |

| | | | | |
|---|---|---|---|---|
| | IGHV4-39 | Downregulated | -4.903 | <0.001 |
| | IGKV1-16 | Downregulated | -4.177 | <0.001 |
| | IGKV3D-20 | Downregulated | -1.299 | <0.001 |
| | IGLV2-8 | Downregulated | -3.87 | <0.001 |
| | IGLV6-57 | Downregulated | -2.307 | <0.001 |
| | ITGB1 | Downregulated | -0.926 | <0.001 |
| | LAPTM5 | Downregulated | -0.832 | <0.001 |
| | LIME1 | Downregulated | -1.377 | <0.001 |
| | MYH9 | Downregulated | -1.047 | <0.001 |
| | NME2 | Downregulated | -1.538 | <0.001 |
| | PCBP2 | Downregulated | -0.809 | <0.001 |
| | SMCHD1 | Downregulated | -1.081 | <0.001 |
| | VIM | Downregulated | -1.251 | <0.001 |
| **B Cells** | | | | |
| | ATP6V0C | Downregulated | -1.346 | <0.001 |
| | BIRC3 | Upregulated | 0.662 | <0.001 |
| | EBLN3 | Upregulated | 0.559 | <0.001 |
| | GPX1 | Upregulated | 1.088 | <0.001 |
| | HLA-DRB5 | Downregulated | -1.275 | <0.001 |
| | HMHA1 | Upregulated | 0.635 | <0.001 |
| | IGHV3-23 | Downregulated | -1.504 | <0.001 |
| | IGHV3-30 | Downregulated | -1.343 | <0.001 |
| | IGKV3-15 | Downregulated | -1.681 | <0.001 |
| | IGKV3-20 | Downregulated | -2.081 | <0.001 |
| | IGLV1-40 | Downregulated | -1.645 | <0.001 |
| | IGLV1-51 | Downregulated | -1.152 | <0.001 |
| | NME2 | Downregulated | -1.533 | <0.001 |
| | PCBP2 | Downregulated | -1.138 | <0.001 |
| | PDE4B | Downregulated | -0.99 | <0.001 |
| | PPDPF | Downregulated | -1.099 | <0.001 |
| | RPL23 | Upregulated | 1.589 | <0.001 |
| | RPS20 | Upregulated | 1.964 | <0.001 |
| | SF1 | Downregulated | -0.952 | <0.001 |
| | SYNGR2 | Downregulated | -0.894 | <0.001 |
| | TNFRSF13C | Upregulated | 0.46 | <0.001 |
| | WHSC1L1 | Upregulated | 0.668 | <0.001 |
| | ZFP36 | Downregulated | -0.954 | <0.001 |
